# Protein homeostasis is maintained by proteasomes containing PSMB9 induced by EEF1A2 upon mitochondrial stress

**DOI:** 10.1101/2021.12.23.473808

**Authors:** Minji Kim, Lukasz Samluk, Tomasz Maciej Stępkowski, Ida Suppanz, Remigiusz Adam Serwa, Agata Kodroń, Bettina Warscheid, Agnieszka Chacinska

## Abstract

Perturbed proteostasis and mitochondrial dysfunction are often associated with age-related diseases such as Alzheimer’s and Parkinson’s diseases. However, the link between them remains incompletely understood. Mitochondrial dysfunction causes proteostasis imbalance, and cells respond to restore proteostasis by increasing proteasome activity and molecular chaperons in yeast and *C. elegans*. Here, we demonstrate the presence of similar responses in humans. Mitochondrial dysfunction upregulates a small heat shock protein HSPB1 and an immunoproteasome subunit PSMB9 leading to an increase in proteasome activity. HSPB1 and PSMB9 are required to prevent protein aggregation upon mitochondrial dysfunction. Moreover, PSMB9 expression is dependent on a translation elongation factor EEF1A2, and PSMB9-containing proteasomes are located near mitochondria, enabling fast local degradation of aberrant proteins. Our findings put a step forward in understanding the stress response triggered by mitochondrial dysfunction, and may be useful for therapeutic strategies to prevent or delay the onset of age-related diseases and attenuate their progression.

## INTRODUCTION

The proteasome is one of the main protein degradation machineries that remove damaged, unfolded and misfolded proteins in eukaryotes. It is composed of two multisubunit subcomplexes: the 20S proteasome which is also called catalytic core particle (CP), and the 19S regulatory particle (RP). The 20S proteasome consists of 4 stacked rings of 28 subunits; 2 rings are composed of 7 different non-catalytic α subunits and 2 rings are composed of 7 different β subunits, three of which are catalytic. The 19S RP is involved in the recognition and processing of ubiquitinated substrates destined for degradation. The 20S proteasome itself can degrade proteins in the ubiquitin and adenosine triphosphate (ATP) independent pathway, but also either one or two 19S RPs can attach to a single 20S proteasome to form the singly capped 26S (RPCP) or the doubly capped 26S proteasome (RP_2_CP), and degrade proteins in the ubiquitin-and ATP-dependent pathway (Collins & Goldberg, 2017, Livneh, Cohen-Kaplan et al., 2016, Pickering & Davies, 2012, Tanaka, 2009). Proteasome activity declines with aging which results in increased protein aggregation - a common feature of age-related diseases, such as neurodegenerative and cardiovascular diseases (Schmidt & Finley, 2014).

Another common feature of age-related diseases is mitochondrial dysfunction. Mitochondria are multifunctional organelles in eukaryotic cells that participate in energy production, metabolism, apoptosis, and cell signaling, and for that reason, their dysfunction can cause various diseases (Chan, 2006, Nunnari & Suomalainen, 2012, Pfanner, Warscheid et al., 2019). Although mitochondria have their own DNA, most mitochondrial proteins are encoded in the nucleus, synthesized as precursor forms in the cytosol and imported into mitochondria by specific transport pathways (Chacinska, Koehler et al., 2009, Neupert & Herrmann, 2007, Wiedemann & Pfanner, 2017). Failure in the import of mitochondrial proteins results in the accumulation of mitochondrial precursor proteins in the cytosol, causing proteotoxic stress. In yeast, mitochondrial proteins are degraded by the proteasome not only under mitochondrial protein import failure but also under physiological conditions, indicating that the proteasome continuously monitors and controls mitochondrial proteins in the cytosol (Bragoszewski, Gornicka et al., 2013). We and another group discovered that impaired mitochondrial protein import triggers the activation of unfolded protein response activated by mistargeting of proteins (UPRam) and mitochondrial precursor overaccumulation stress (mPOS) to modulate protein translation and protein degradation in yeast (Wang & Chen, 2015, Wrobel, Topf et al., 2015). In both yeast and human cells, mitochondrial defects cause a decrease in global translation that may act to reduce the burden of newly synthesized proteins, otherwise prone to aggregate (Samluk, Urbanska et al., 2019, Wrobel et al., 2015). In addition, mitochondrial protein import stress enhances proteasome activity in yeast (Wrobel et al., 2015) and similarly, mitochondrial defects increase proteasome activity and extend life span in *Caenorhabditis elegans* (*C.elegans*) (Sladowska, Turek et al., 2021). Yet, it remains unclear whether and how the proteasome is activated in response to mitochondrial dysfunction in humans.

Heat shock proteins (HSPs), a large family of molecular chaperones, constitute another layer of protein quality control mechanisms acting to protect proteins from aggregation (Balchin, Hayer-Hartl et al., 2016, Glover & Lindquist, 1998). The mitochondrial unfolded protein response (UPRmt), one of the best-characterized stress responses mainly studied in *C.elegans*, is activated by mitochondrial proteotoxic stress leading to the induction of the transcription of genes encoding proteases and HSPs located in the mitochondrial matrix through mitochondria-to-nucleus communication (Labbadia, Brielmann et al., 2017, Nargund, Pellegrino et al., 2012, Yoneda, Benedetti et al., 2004). However, there is limited research focused on how cytosolic HSPs are regulated to protect the cell from mitochondrial proteotoxic stress. It was reported that mitochondrial import stress upregulates the gene expression of HSPs and proteasome subunits and downregulates the gene expression of cytosolic translation components in yeast (Boos, Kramer et al., 2019). We recently also found in yeast that mitochondrial precursor proteins, which are accumulated in the cytosol upon mitochondrial protein import failure, lead to the upregulation of specific HSPs to cope with the burden of mislocalized mitochondrial proteins and prevent their aggregation with non-mitochondrial proteins in the cytosol (Nowicka, Chroscicki et al., 2021).

In the present study, we have explored stress responses that are activated upon mitochondrial dysfunction and act to protect the cell against proteotoxicity in humans. We have observed selective upregulation of a small HSP family member HSPB1 and an increase in the proteasome activity. We identified an inducible immunoproteasome β subunit PSMB9 as a key factor of proteasome activity enhancement upon mitochondrial dysfunction. Moreover, we have found that upregulation of the translation elongation factor EEF1A2, and it was required for PSMB9 induction under mitochondrial stress. Importantly, depleting HSPB1, PSMB9 or EEF1A2 accelerated protein aggregation, reflecting their roles to safeguard cells against mitochondrial dysfunction-mediated proteotoxicity.

## RESULTS

### Mitochondrial complex I deficiency upregulates HSPB1 and HSPH1

To study cellular stress responses activated upon mitochondrial dysfunction in humans, we took advantage of two different mitochondrial complex I-deficient human cells, TALEN-based *NDUFA11* knockout (KO) and *NDUFA13* KO human embryonic kidney 293T (HEK293T) cells. Mitochondrial complex I is the first and the largest enzyme of the mitochondrial respiratory chain composed of 45 subunits in humans, and its deficiency is the most common enzyme defect in mitochondrial disorders (Mimaki, Wang et al., 2012). NDUFA11 and NDUFA13 are accessary subunits of mitochondrial complex I, and their deficiencies in HEK293T cells cause a significant loss of complex I assembly (Stroud, Surgenor et al., 2016). We previously found that *NDUFA11* KO has higher reactive oxygen species (ROS) production and lower ATP generation compared with wild-type (WT) HEK293T cells (Samluk et al., 2019). In this study, we first investigated the global transcriptomic changes that were triggered by NDUFA11 and NDUFA13 deficiency. In total 13,502 genes were detected, with 651 genes significantly upregulated (log_2_ fold change > 1, q-value < 0.05), and 318 genes significantly downregulated (log_2_ fold change < −0.75, q-value < 0.05) in both *NDUFA11* KO and *NDUFA13* KO compared to WT HEK293T cells. With these genes, we performed the Reactome pathway enrichment analysis (Supplementary Fig. 1). The analysis showed that the expression of genes involved in extracellular matrix (ECM) organization was significantly upregulated in mitochondrial complex I-deficient HEK293T cells compared to WT HEK293T cells, which is important to return to their normal physical equilibrium under stress conditions (Bonnans, Chou et al., 2014). The expression of genes involved in translation was significantly downregulated in mitochondrial complex I-deficient HEK293T cells compared to WT HEK293T cells, indicating a cellular attempt to limit protein burden upon mitochondrial dysfunction. We then assessed differences in gene expression of HSPs upon mitochondrial dysfunction. The genes of small HSP and HSP110 families had an upregulated tendency, while those of HSP10, HSP60, HSP70 and HSP90 families had a downregulated tendency in mitochondrial complex I-deficient HEK293T cells compared to WT in RNA-sequencing (RNA-seq) analysis (Fig. 1a). qRT-PCR validation experiments for 6 selected HSPs were consistent with the RNA-seq results (Fig. 1b), so was western blotting validation (Fig. 1c). These data show that HSPB1 and HSPH1 were markedly upregulated at the mRNA and protein level, whereas levels of HSP1A1/HSP1AB and HSP90 were downregulated or unchanged upon mitochondrial dysfunction. *NDUFA11* KO resulted in the decrease of NDUFA13 protein expression, and vice versa, as previously reported (Stroud et al., 2016). These results suggest that selectively upregulated HSPs such as HSPB1 could play an important role in protecting cells from protein aggregation under mitochondrial stress conditions.

**Figure 1.**
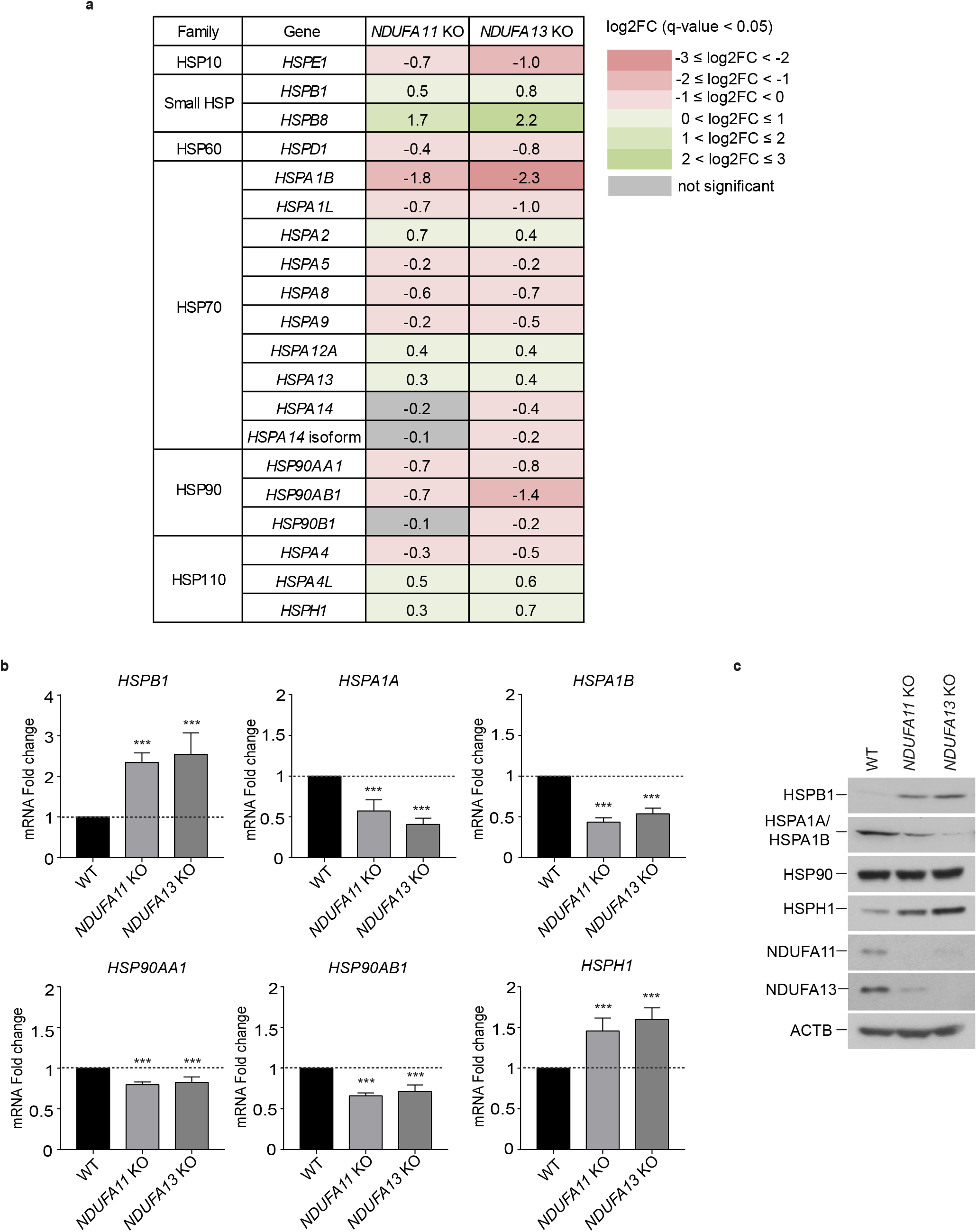
small HSP and HSP110 families have a upregulated tendency under mitochondrial stress. (**a)** RNA-seq analysis of HSPs gene expression log2 fold changes (log2FC) in *NDUFA11* KO and *NDUFA13* KO compared to WT HEK293T cells. Up- and down-regulated genes (q-value < 0.05) are shown in green and pink, respectively. The intensity of the color shades depends on the level of expression change. Gray indicates genes with not statistically significant expression changes. (**b)** mRNA expression patterns of selected transcripts validated by RT-qPCR. The mRNA levels are presented as fold changes relative to WT. Data shown are mean ± SD. ***, p < 0.001 from an ordinary one-way ANOVA with Dunnett’s multiple comparisons test using GraphPad Prism. (**c)** Western blot analysis of HSP expression performed in whole cell lysates of *NDUFA11* KO, *NDUFA13* KO and WT HEK293T cells. ACTB was used as a loading control. Data shown are representative of 3 independent experiments.

### 20S proteasome subunits and HSPB1 are specifically enriched in protein aggregates isolated from *NDUFA11* KO cells

To examine whether the upregulation of HSPB1 and HSPH1 is associated with protein aggregation (Balchin et al., 2016, Webster, Darling et al., 2019, Yamagishi, Ishihara et al., 2003), we enriched insoluble proteins from total lysates of *NDUFA11* KO and WT HEK293T cells by two subsequent centrifugation steps using 1,000 × g and 125,000 × g spins following previously established protocols (Bender, Lewrenz et al., 2011, Nowicka et al., 2021, Walther, Kasturi et al., 2015) and performed mass spectrometry (MS)-based proteomics analysis. Notably, 20S proteasome subunits (red dots) and specific HSPs (blue dots) including HSPB1 were more abundant in aggregates isolated from *NDUFA11* KO compared to those from WT HEK293T cells, whereas 19S proteasome subunits (orange dots) did not show any difference in abundance in aggregates enriched between *NDUFA11* KO *versus* WT HEK293T cells (Fig. 2a). Next, we sought to evaluate the effect of aggregation-prone proteins on HSPB1 induction. To do so, we transfected cells with plasmids encoding enhanced green fluorescent protein (EGFP)-tagged *MAPT (*also known as *tau*) (Hoover, Reed et al., 2010), because HSPB1 has been shown to be involved in the regulation of MAPT aggregation in vitro and in vivo (Abisambra, Blair et al., 2010, Baughman, Clouser et al., 2018). Co-immunoprecipitation assays revealed more interaction of MAPT with both HSPB1 and HSPH1 in *NDUFA11* KO and *NDUFA13* KO compared with WT HEK293T cells (Fig. 2b). MS-based proteomics analysis of aggregates from total lysates of *NDUFA11* KO and WT HEK293T cells expressing EGFP-tagged MAPT showed still more abundant HSPB1 in aggregates of *NDUFA11* KO compared to those of WT HEK293T cells. In contrast, MAPT and proteasome subunits did not show any difference in aggregates between *NDUFA11* KO and WT HEK293T cells that might be due to solubility limitation during sample preparation, or a technical barrier of mass spectrometry having difficulty to detect insoluble proteins (Fig. 2c). No changes in the mRNA and protein expression levels of HSPB1 were observed in either mitochondrial complex I-deficient or WT HEK293T cells by MAPT expression (Fig. 2d, e). We also tested whether *HSPB1* mRNA expression is regulated by heat shock factor (HSF1), a well-known master regulator of the heat shock response (Anckar & Sistonen, 2011). There was no change in the abundance of *HSPB1* mRNA, rather it showed a slightly increasing tendency in both mitochondrial complex I-deficient and WT HEK293T cells after *HSF1* knockdown, therefore, the *HSPB1* upregulation was independent of HSF1 upon mitochondrial dysfunction (Supplementary Fig. 2). To conclude, our results indicate that mitochondrial dysfunction rather than an exogenous aggregation-prone protein is a driving force for HSPB1 induction, and the induced HSPB1 is associated with protein aggregation.

**Figure 2.**
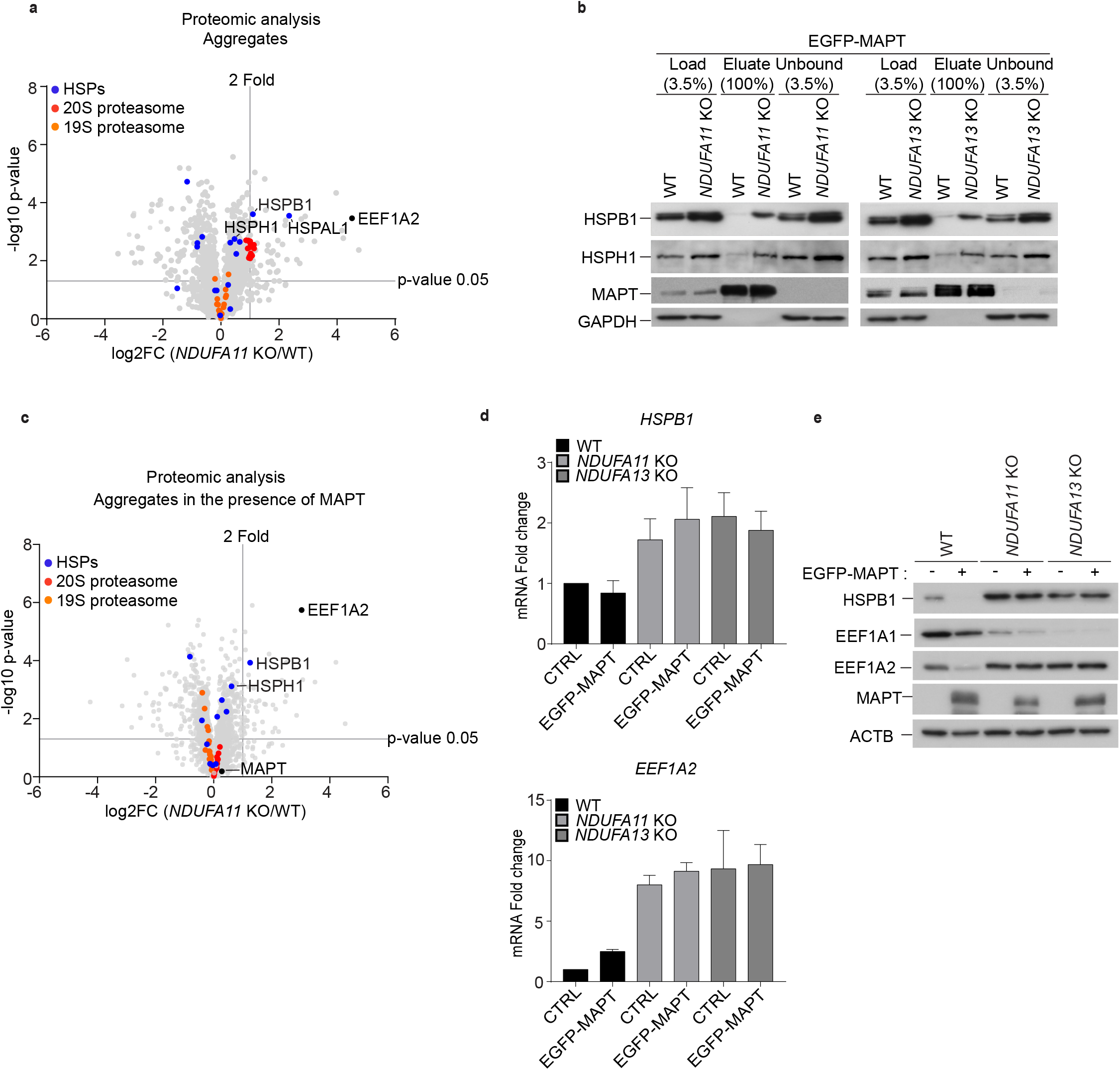
20S proteasome subunits and HSPB1 are enriched in protein aggregates isolated from *NDUFA11* KO HEK293T cells. (**a**, **c)** Volcano plots displaying the log2 fold change (log2FC, x axis) against the t test-derived −log10 statistical p-value (y axis) for all proteins detected in aggregate fractions of *NDUFA11* KO and WT HEK293T cells by LC-MS/MS analysis in the absence (**a**) and in the presence (**c**) of EGFP-MAPT, respectively. 20S proteasome subunits, 19S proteasome subunits and HSPs are indicated by red, orange and blue dots, respectively. EEF1A2 and MAPT are indicated by black dots. (**b**) Immunoprecipitation followed by western blot analysis of EGFP-*MAPT* transfected cell lysates performed with anti-MAPT antibody. Load and unbound: 3.5%, Eluate: 100%. GAPDH was used as a loading control. (**d**) mRNA expression levels of *HSPB1* and *EEF1A2* examined by RT-qPCR analysis. The mRNA levels are presented as fold changes relative to WT (n=2). (**e**) Western blot analysis of HSPB1 and EEF1A1/2 expression performed in whole cell lysates of *NDUFA11* KO, *NDUFA13* KO and WT HEK293T cells. ACTB was used as a loading control. Data shown are representative of 3 independent experiments.

### Mitochondrial complex I deficiency increases *PSMB9* mRNA level, and enhances proteasome activity

Given that the 20S proteasome was more abundant in aggregates of *NDUFA11* KO compared to WT (Fig. 2a), we assessed changes in gene expression of proteasome subunits upon mitochondrial dysfunction. Intriguingly, most genes of proteasome subunits were downregulated but an inducible 20S immunoproteasome subunit, *PSMB9,* and a 19S proteasome subunit, *PSMD10* were upregulated in *NDUFA11* KO and *NDUFA13* KO compared to WT HEK293T cells as shown by RNA-seq analysis (Fig. 3a). The catalytic β subunits PSMB8, PSMB9 and PSMB10 were identified as interferon γ-inducible subunits in higher eukaryotes that are incorporated into a specialized form of the 20S proteasome, called the immunoproteasome, by replacement of the constitutive 20S proteasome catalytic β subunits PSMB5, PSMB6 and PSMB7, respectively (Schmidt & Finley, 2014, Tanaka, 2009). qRT-PCR validation for *PSMB5*, *PSMB6*, *PSMB7* and *PSMB9* showed similar expression patterns with the RNA-seq results. Although *PSMB8* and *PSMB10* were not detected in RNA-seq analysis, they were detected in qRT-PCR analysis. Nevertheless, *PSMB8* and *PSMB10* both remained unchanged, while *PSMB9* was upregulated in *NDUFA11* KO and *NDUFA13* KO compared to WT HEK293T cells (Fig. 3b). To investigate the impact of mitochondrial dysfunction on proteasome activity, chymotrypsin-like and caspase-like proteasome activities were measured using fluorigenic peptide substrates (Fig. 3c). Their levels were significantly higher in *NDUFA11* KO and *NDUFA13* KO compared to WT HEK293T cells, similar to previous observations in yeast and *C.elegans* upon mitochondrial dysfunction (Sladowska et al., 2021, Wrobel et al., 2015), demonstrating that mitochondrial defects trigger cellular stress responses such as UPRam also in humans. We next examined whether mitochondrial stress alters proteasome assembly using native polyacrylamide gel electrophoresis (PAGE). The 20S proteasome was more abundant in *NDUFA11* KO and *NDUFA13* KO compared to WT HEK293T cells, whereas the 26S proteasome remained unchanged (Fig. 3d). There was no difference between mitochondrial complex I-deficient and WT HEK293T cells in the protein expression levels of α subunits of the 20S proteasome and a 19S proteasome subunit (Fig. 3e). Since the 20S proteasome participates in the ubiquitin-independent degradation pathway, we examined whether global ubiquitination is affected by mitochondrial dysfunction. Proteasome inhibition by MG132 treatment elevated global protein ubiquitination, however, no detectable difference was observed between mitochondrial complex I-deficient and WT HEK293T cells, indicating that the increased proteasome activity in mitochondrial complex I-deficient HEK293T cells functions ubiquitin-independently (Supplementary Fig. 3).

**Figure 3.**
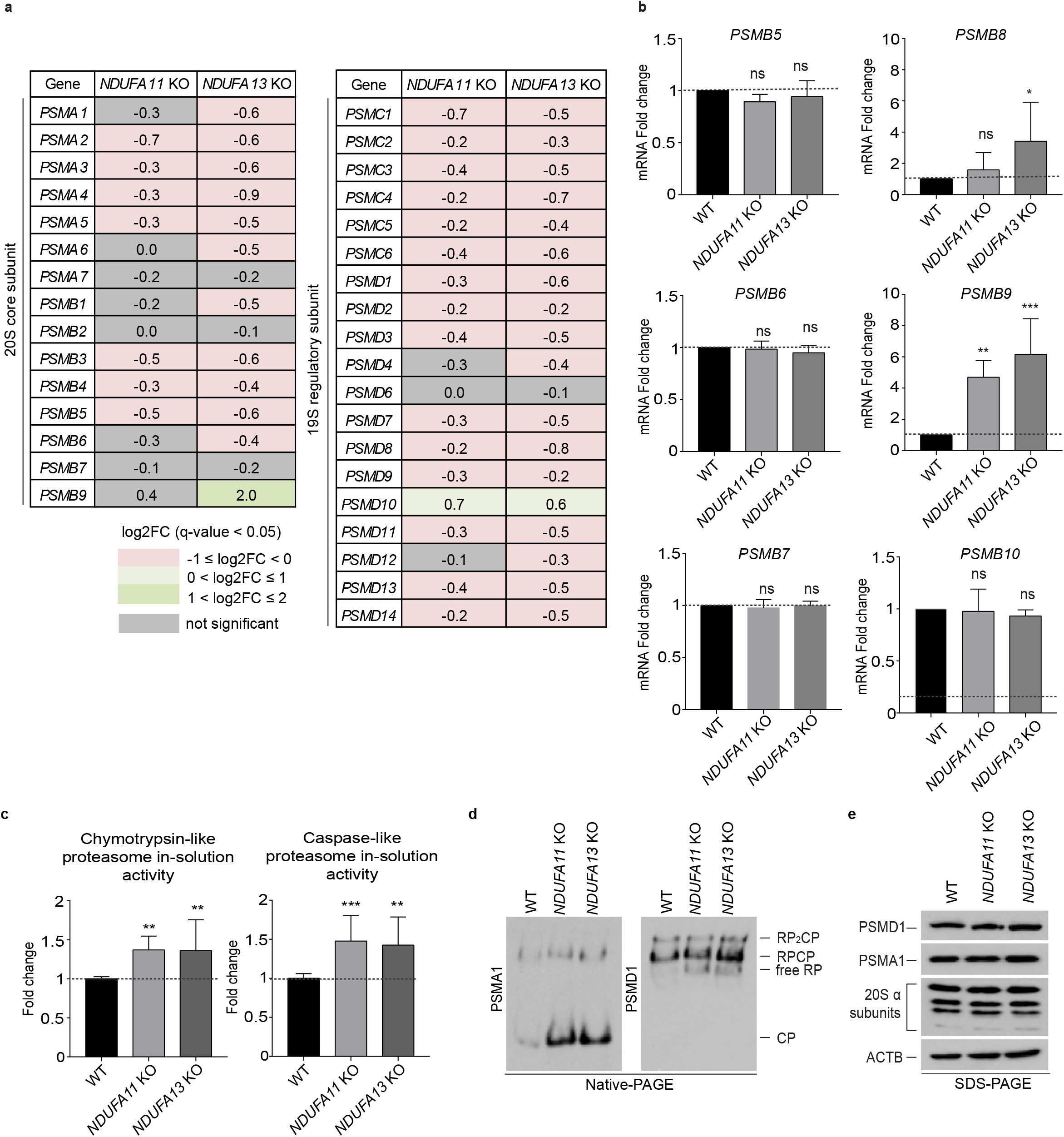
Mitochondrial complex I deficiency increases PSMB9 mRNA level and enhances proteasome activity. (**a**) RNA-seq analysis of 20S (left panel) and 19S (right panel) proteasome components gene expression log2 fold changes (log2FC) in mitochondrial complex I-deficient HEK293T cells compared to WT HEK293T cells. Up- and down-regulated genes (q-value < 0.05) are shown in green and pink, respectively. The intensity of the color shades depends on the level of expression change. Gray indicates genes with not statistically significant expression changes. (**b**) mRNA expression patterns of selected transcripts validated by RT-qPCR. The mRNA levels are presented as fold changes relative to WT. (**c**) Chymotrypsin-like and caspase-like proteasome activities in cell lysates presented as fold changes relative to WT. Data shown are mean ± SD (n=5). *p < 0.05, ** p < 0.01, *** p < 0.001; ns, not significant from an ordinary one-way ANOVA with Dunnett’s multiple comparisons test using GraphPad Prism. (**d**) Proteasome species in *NDUFA11* KO, *NDUFA13* KO and WT HEK293T cell extracts resolved by electrophoresis in 4.5% native gel followed by western blot analysis detecting a 20S proteasome subunit PSMA1 and a 19S proteasome subunit PSMD1 to characterize 26S (RP_2_CP, doubly capped 26S; RP_1_CP, singly capped 26S) and 20S (CP, core particle) proteasomes. (**e**) Western blot analysis of proteasome subunit expression performed in whole cell lysates of mitochondrial complex I-deficient and WT HEK293T cells. ACTB was used as a loading control. Data shown are representative of 3 independent experiments.

### Mitochondrial stress-induced PSMB9 is responsible for proteasome activity enhancement

To characterize the proteasome composition in detail, plasmids expressing FLAG-tagged proteasome subunit alpha type-5 (PSMA5) were generated and expressed in *NDUFA11* KO, *NDUFA13* KO and WT HEK293T cells, and proteasomes were purified by anti-FLAG immunoprecipitation and analyzed using a quantitative MS-based approach (Fig. 4a). As a result, PSMB9 was found to be more abundant in proteasomes purified from *NDUFA11* KO compared with those from WT HEK293T cells (Fig. 4b, upper panel). PSMB9 was also more abundant in proteasomes purified from *NDUFA13* KO than those from WT HEK293T cells, although it wasn’t statistically significant (Fig. 4b, lower panel). Of note, HSPB1 was increased in abundance in proteasome fractions purified from both *NDUFA11* KO and *NDUFA13* KO compared with those from WT HEK293T cells (Fig. 4b). This finding suggests that HSPB1 might assist proteasomes in efficient protein degradation upon mitochondrial dysfunction. In line with this, western blot analysis showed an induction of PSMB9 protein expression in *NDUFA11* KO and *NDUFA13* KO HEK293T cells (Fig. 4c, Supplementary Fig. 4a). In contrast, there was no difference in the levels of PSMB5, PSMB6 and PSMB8 between mitochondrial complex I-deficient and WT HEK293T cells. Native PAGE analysis further demonstrated the incorporation of PSMB9 into proteasomes in *NDUFA11* KO and *NDUFA13* KO HEK293T cells (Fig. 4d), along with an increase in PSMB9-specific proteasome activity in *NDUFA11* KO and *NDUFA13* KO compared to WT HEK293T cells as measured using fluorigenic peptide substrate specific for PSMB9 (Fig. 4e). Moreover, *PSMB9* depletion using small interfering RNA (siRNA) decreased chymotrypsin-like and caspase-like proteasome activities in *NDUFA11* KO and *NDUFA13* KO HEK293T cells (Fig. 4f).

**Figure 4.**
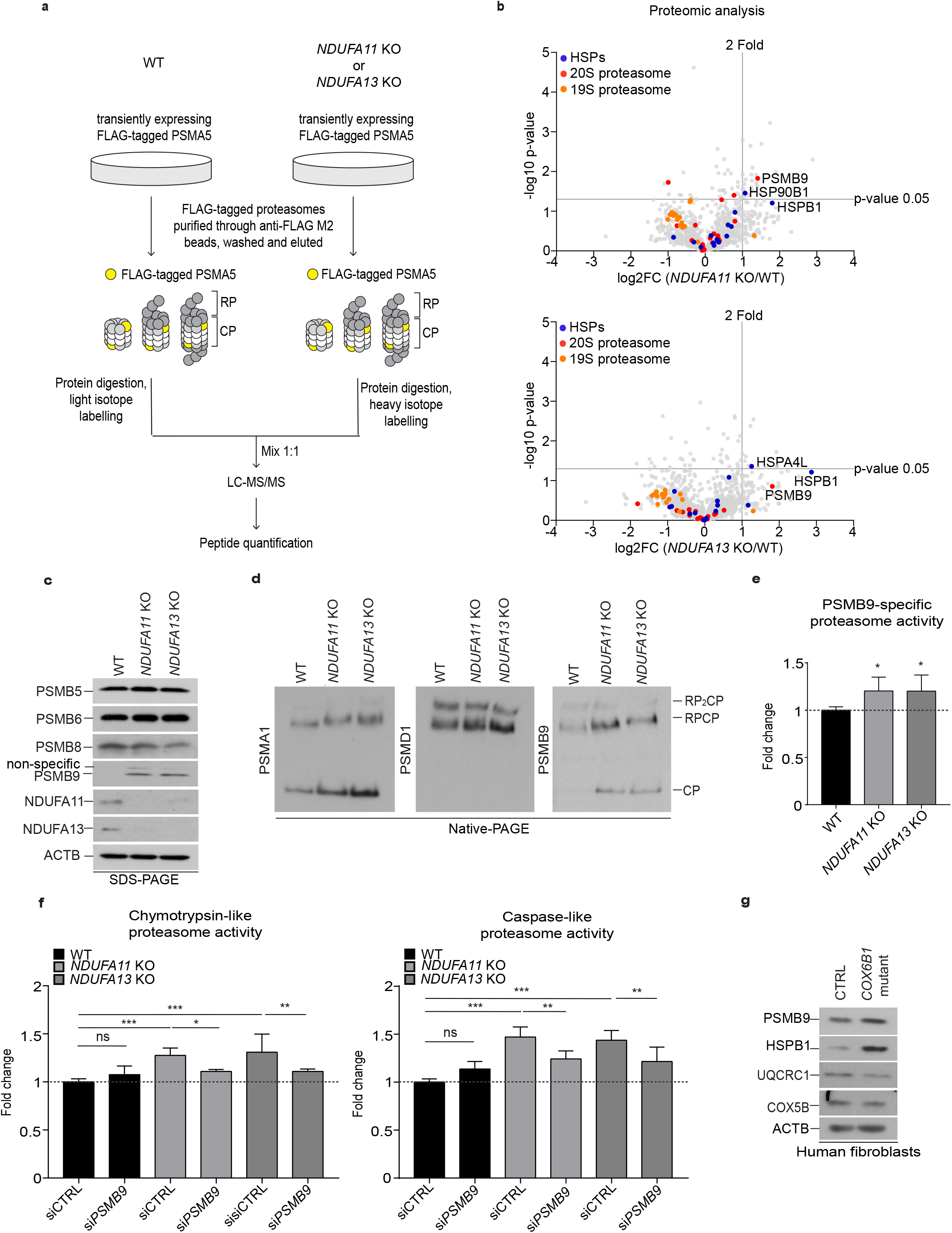
PSMB9 is more abundant in proteasomes purified from mitochondrial complex I-deficient cells compared to those of WT. (**a**) Workflow of the affinity purification of FLAG-PSMA5 tagged proteasomes using anti-FLAG affinity gels for LC-MS/MS analysis. (**b**) Volcano plots displaying the log2 fold change (log2FC, x axis) against the t test-derived −log10 statistical p-value (y axis) for all proteins identified in purified proteasomes of *NDUFA11* KO versus WT (upper panel) or *NDUFA13* KO versus WT (lower panel) HEK293T cells by LC-MS/MS analysis. 20S proteasome subunits, 19S proteasome subunits and HSPs are indicated by red, orange and blue dots, respectively. (**c**) Western blot analysis of the expression of proteasome β subunits performed in whole cell lysates of mitochondrial complex I-deficient and WT HEK293T cells. ACTB was used as a loading control. (**d**) Proteasome species in mitochondrial complex I-deficient and WT HEK293T cell extracts resolved by electrophoresis in 4.5% native gel followed by western blot analysis using a 20S proteasome subunit PSMA1, 20S immunoproteasome subunit PSMB9 and a 19S proteasome subunit PSMD1 to characterize 26S (RP_2_CP, doubly capped 26S; RP_1_CP, singly capped 26S) and 20S (CP, core particle) proteasomes. Data shown are representative of 3 independent experiments. (**e**) PSMB9-specific proteasome activity in cell lysates presented as fold change relative to WT. Data shown are mean ± SD. *p < 0.05 from an ordinary one-way ANOVA with Dunnett’s multiple comparisons test using GraphPad Prism. (**f**) Chymotrypsin-like and caspase-like proteasome activities in cell lysates presented as fold changes relative to WT 72 h after transfection with *PSMB9* siRNA or control siRNA. Data shown are mean ± SD. *p < 0.05, ** p < 0.01, *** p < 0.001; ns, not significant from an ordinary one-way ANOVA with Tukey’s multiple comparisons test using GraphPad Prism. (**g**) Western blot analysis performed in whole cell lysates of *COX6B1*-mutant and control fibroblasts. ACTB was used as a loading control. Data shown are representative of 3 independent experiments.

A rescue experiment by overexpression of *NDUFA11* in *NDUFA11* KO HEK293T cells using FLAG-tagged *NDUFA11* plasmid reversed the protein change of HSPB1 and PSMB9, confirming that their induction was elicited by the absence of *NDUFA11* (Supplementary Fig. 4b). We further tested whether mitochondrial stress triggered by chemical treatments in WT cells induces HSPB1 and PSMB9 expression as we observed in mitochondrial complex I-deficient HEK293T cells. To this end, WT HEK293T cells were treated with rotenone (mitochondrial complex I inhibitor), menadione (mitochondrial ROS inducer), and carbonyl cyanide m-chlorophenylhydrazone (CCCP; mitochondrial depolarizer) for 2 and 24 hours (h) followed by western blot analysis. None of treatments induced HSPB1 and PSMB9 protein expression (Supplementary Fig. 4c). In addition, we reanalyzed published transcriptomic and proteomic data of HeLa cells treated by actinonin (mitochondrial protein synthesis inhibitor), carbonylcyanide-4-(trifluoromethoxy)phenylhydrazone (FCCP; mitochondrial depolarizer), and MitoBloCK-6 (MB; mitochondrial import inhibitor of both CHCHD4 and TIM22 import pathways) (Quiros, Prado et al., 2017). Interestingly, PSMB9 gene and protein expression was upregulated in MB-treated HeLa cells (Supplementary Fig. 4d). These findings suggest that HSPB1 and PSMB9 induction is led by *NDUFA11* depletion, and mitochondrial import stress may be associated with PSMB9 upregulation.

To investigate whether our observations from mitochondrial complex I-deficient HEK293T cells are relevant to other mitochondrial disease models, we used human fibroblasts from a patient with a mutation in a nucleus-encoded mitochondrial respiratory chain complex IV subunit COX6B1, presenting an early-onset encephalopathy with leukodystrophy, myopathy and growth retardation associated with Cytochrome c oxidase (COX) deficiency (Massa, Fernandez-Vizarra et al., 2008). *COX6B1*-mutant fibroblasts had lower protein expression of UQCRC1, a subunit of mitochondrial complex III, and COX5B, a subunit of mitochondrial complex IV, indicating mitochondrial defects. Importantly, elevated protein expression of HSPB1 and PSMB9 was observed in *COX6B1*-mutant fibroblasts, suggesting that a similar stress response is activated by COX deficiency in fibroblasts (Fig. 4g).

To conclude, our data indicate that induced PSMB9 is responsible for enhancement of proteasome activity upon mitochondrial stress.

### Mitochondrial stress-induced EEF1A2 regulates PSMB9 expression

Translation is the process of converting mRNA to an amino acid chain and it occurs in the three steps: initiation, elongation, and termination. Aminoacylated-tRNAs are delivered to the ribosome by the translation elongation factor EEF1A. The role of EEF1A is guanosine triphosphate (GTP)-dependent, and this process is facilitated by a GTP-exchange factor called EEF1B. In mammals, there are two isoforms of EEF1A, EEF1A1 and EEF1A2 (Knudsen, Frydenberg et al., 1993, Mateyak & Kinzy, 2010). Unexpectedly, we observed that EEF1A2 is strongly enriched in aggregates of *NDUFA11* KO compared to WT HEK293T cells (Fig. 2a, c). Thus, we assessed changes in gene expression of translation factors including EEF1A2 upon mitochondrial dysfunction. Most genes of translation factors were downregulated but only three genes were upregulated, and *EEF1A2* was the most upregulated gene among them in *NDUFA11* KO and *NDUFA13* KO compared to WT HEK293T cells in RNA-seq analysis (Fig. 5a). The downregulation of *EEF1A1* and upregulation of its isoform *EEF1A2* were confirmed by qRT-PCR and western blot data (Fig. 5b, c). Similar to HSPB1, no change in the mRNA and protein expression of EEF1A2 was observed in mitochondrial complex I-deficient or WT HEK293T by MAPT expression. There was a slight increase in *EEF1A2* mRNA expression by MAPT expression in WT, however, its protein expression was rather decreased indicating that EEF1A2 was upregulated in response to mitochondrial dysfunction rather than MAPT (see Fig. 2d, e). EEF1A1 was decreased by rotenone and CCCP treatment, while EEF1A2 remained unchanged by chemical mitochondrial stressors (Supplementary Fig. 4b). These data led us to hypothesize that mitochondrial complex I deficiency induces EEF1A2 expression, and it participates in selective translation of specific mRNA such as *HSPB1* and *PSMB9*.

**Figure 5.**
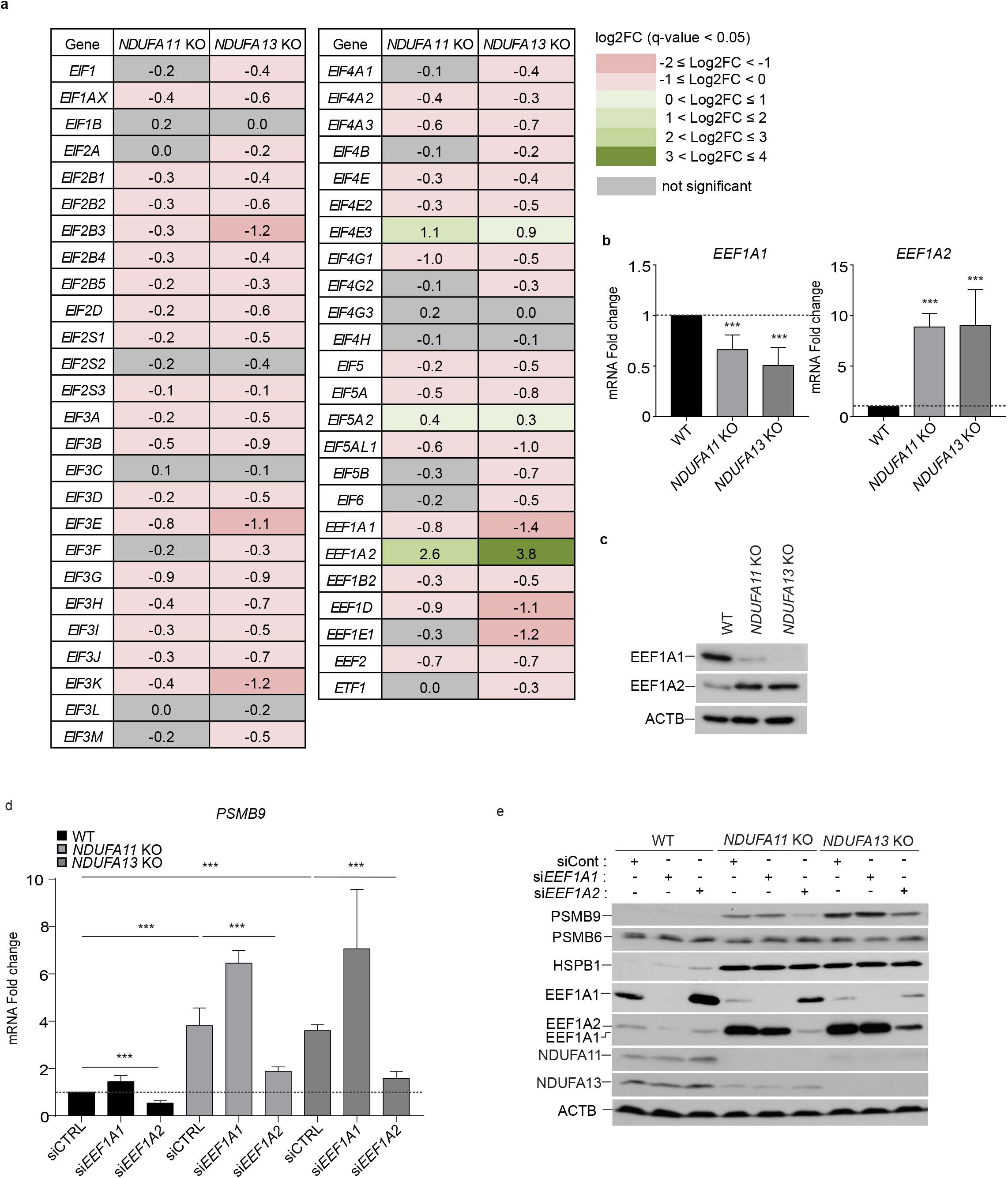
EEF1A2 are induced under mitochondrial stress. (**a**) RNA-seq analysis of translation factor gene expression log2 fold changes (log2FC) in mitochondrial complex I-deficient cells compared to WT HEK293T cells. Up- and down-regulated genes (q-value < 0.05) are shown in green and pink, respectively. The intensity of the color shades depends on the level of expression change. Gray indicates genes with not statistically significant expression changes. (**b**) mRNA expression patterns of *EEF1A1* and *EEF1A2* by RT-qPCR. The mRNA levels are presented as fold changes relative to WT. Data shown are mean ± SD. ***p < 0.001 from an ordinary one-way ANOVA with Dunnett’s multiple comparisons test using GraphPad Prism. (**c**) Western blot analysis of EEF1A1 and EEF1A2 performed in whole cell lysates of WT and mitochondrial complex I-deficient HEK293T cells. (**d**, **e**) Mitochondrial complex I-deficient HEK293T cells and WT HEK293T cells were transfected with *EEF1A1*, *EEF1A2* or control siRNAs for 72 h. (**d**) *PSMB9* mRNA expression pattern examined by RT-qPCR in WT and mitochondrial complex I-deficient HEK293T cells. The mRNA levels are presented as fold changes relative to WT transfected with control siRNA. Data shown are mean ± SD. ***p < 0.001 from two-tailed, unpaired t-test using GraphPad Prism. (**e**) Western blot analysis performed in whole cell lysates of mitochondrial complex I-deficient HEK293T cells and WT HEK293T cells. ACTB was used as a loading control. Data shown are representative of 3 independent experiments.

To investigate the role of EEF1A2 on HSPB1 and PSMB9 protein expression, we used siRNAs to deplete *EEF1A2* in mitochondrial complex I-deficient and WT HEK293T cells. *EEF1A1* was also depleted to compare the effects of the two different translation elongation factors on HSPB1 and PSMB9 protein expression. Surprisingly, *EEF1A2* knockdown significantly reduced not only protein expression but also mRNA expression of PSMB9 (Fig. 5d, e). *EEF1A1* knockdown increased *PSMB9* mRNA expression, however, PSMB9 protein expression was not much affected (Fig 5d, e). Neither *EEF1A1* nor *EEF1A2* knockdown changed the mRNA and protein expression of *PSMB6* and *HSPB1* (Fig. 5e, Supplementary Fig. 5a). Furthermore, we found evidence of compensatory mechanisms between EEF1A1 and EEF1A2. *EEF1A1* knockdown markedly increased *EEF1A2* mRNA expression, whereas its protein expression was unaffected in both mitochondrial complex I-deficient and WT HEK293T cells. *EEF1A2* knockdown did not change mRNA expression of *EEF1A1*, whereas EEF1A1 protein expression was increased in both mitochondrial complex I-deficient and WT HEK293T cells, so that even it was recognized by anti-EEF1A2 antibody in WT HEK293T cells due to their high amino acid sequence similarity, indicating that cells have complicated regulations in the process linking gene expression to protein synthesis of EEF1A1 and EEF1A2 (Fig. 5e, Supplementary Fig. 5a).

To test the possibility that induced EEF1A2 stabilizes *PSMB9* mRNA, *PSMB9* mRNA stability was examined after a transcription inhibitor actinomycin D (ActD) treatment up to 24 h. *PSMB9* mRNA stability was not higher in mitochondrial complex I-deficient HEK293T cells compared to WT HEK293T cells (Supplementary Fig. 5b), therefore, EEF1A2 regulates the transcription and translation of PSMB9 rather than its mRNA stability.

### PSMB9 and HSPB1 are required to maintain proteostasis upon mitochondrial dysfunction

Mitochondrial dysfunction has been reported to promote protein aggregate formation in yeast, *C.elegans* and human cells (Biel, Aryal et al., 2020, Nowicka et al., 2021). Hence, we investigated whether mitochondrial complex I deficiency triggers protein aggregation, and whether PSMB9 and HSPB1 have physiological functions on the accumulation of protein aggregates. To elucidate the effect of PSMB9 and HSPB1 in protein aggregation, *PSMB9* or *HSPB1* was depleted using siRNAs (Supplementary Fig. 6a), and protein aggregates in *NDUFA11* KO and WT HEK293T cells were stained with ProteoStat® dye, and monitored by confocal microscopy. *NDUFA11* KO had more protein aggregates than WT HEK293T cells, and *PSMB9* or *HSPB1* single knockdown significantly increased protein aggregation in *NDUFA11* KO HEK293T cells. *PSMB9* and *HSPB1* double knockdown showed a similar deleterious effect with single depletion of *PSMB9* or *HSPB1*, implying their collaboration act in the same pathway under mitochondrial stress against protein aggregation. Conversely, no increase of protein aggregates was observed in WT HEK293T cells (Fig. 6a, b). We also examined the effect of *PSMB6* silencing on proteasome activities and protein aggregation. *PSMB6* knockdown significantly decreased chymotrypsin-like and caspase-like proteasome activities in both mitochondrial complex I-deficient and WT HEK293T cells (Supplementary Fig. 6b, c). However, unlike *PSMB9* knockdown, *PSMB6* knockdown did not increase protein aggregation in both *NDUFA11* KO and WT HEK293T cells, emphasizing the importance of PSMB9 for preventing protein aggregation upon mitochondrial dysfunction (Supplementary Fig. 6d, e). We extended the observation of protein aggregates to cells after *EEF1A2* or *HSF1* knockdown. *EEF1A2* knockdown remarkably promoted protein aggregate formation in *NDUFA11* KO HEK293T cells whereas *HSF1* knockdown did not in *NDUFA11* KO HEK293T cells. Neither *EEF1A2* nor *HSF1* knockdown changed protein aggregate formation in WT HEK293T cells (Fig. 6c, d). To further study the role of PSMB9 and HSPB1 in pathological protein aggregation, we observed MAPT aggregation in EGFP-MAPT expressing *NDUFA11* KO and WT HEK293T cells by confocal microscopy. Similar to ProteoStat® dye, more strong EGFP fluorescence intensities indicating more MAPT aggregation were observed in *NDUFA11* KO than in WT HEK293T cells. *PSMB9* or *HSPB1* single knockdown, and *PSMB9* and *HSPB1* double knockdown significantly increased MAPT aggregation in *NDUFA11* KO HEK293T cells. In contrast, no increase of MAPT aggregates was observed in WT HEK293T cells (Fig. 6e, f). These findings demonstrate that PSMB9 and HSPB1 prevent protein aggregation upon mitochondrial dysfunction, and support that EEF1A2, the upstream regulator of PSMB9, is an important factor to maintain a balanced proteome under mitochondrial stress.

**Figure 6.**
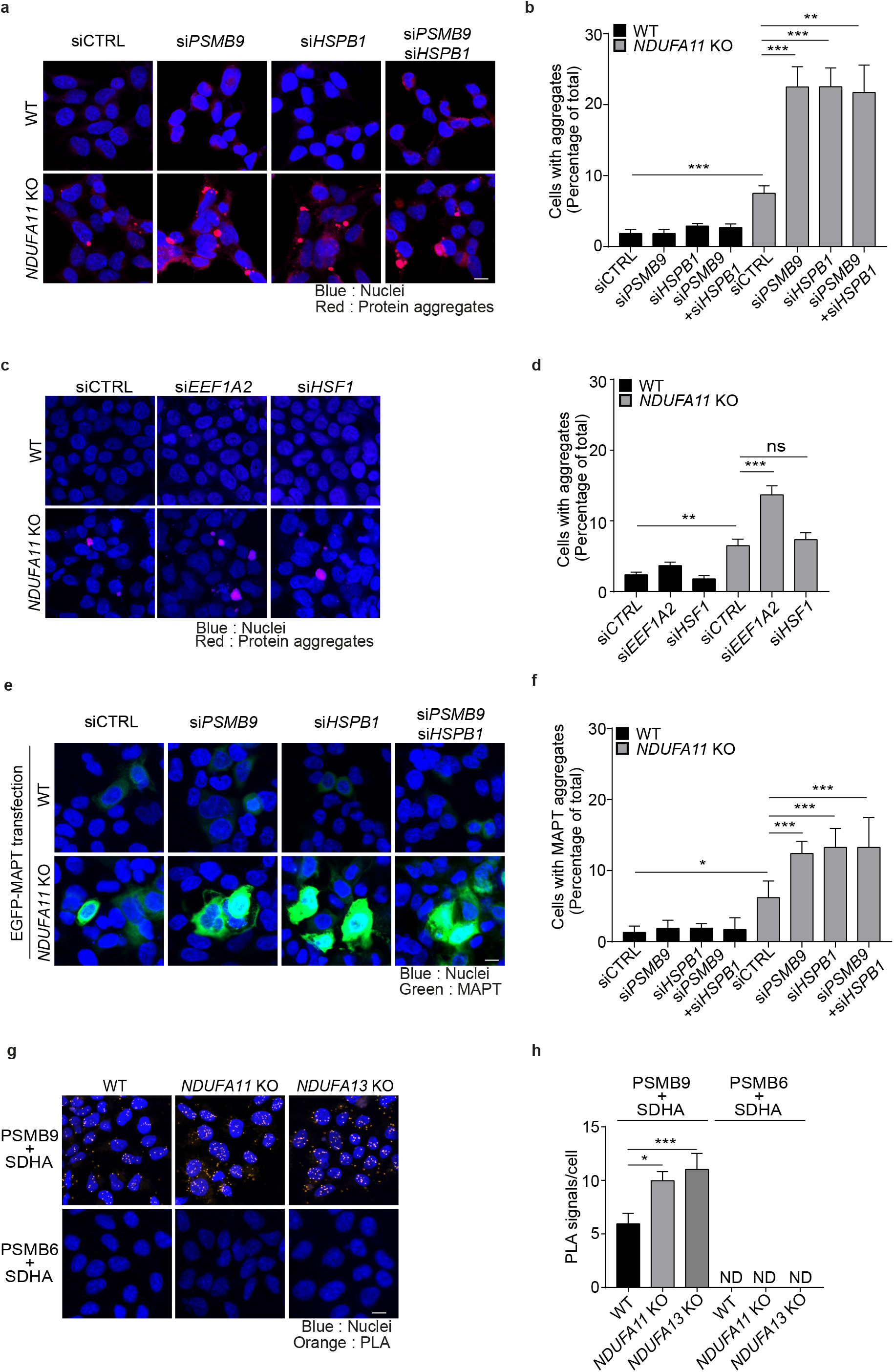
PSMB9 and HSPB1 are required to maintain proteostasis upon mitochondrial dysfunction. (**a**, **b**) *NDUFA11* KO and WT HEK293T cells were transfected with *PSMB9*, *HSPB1*, both *PSMB9* and *HSPB1* or control siRNAs for 72 h. (**a**) Images of protein aggregates stained with ProteoStat. (**b**) Quantification of the percentage of cells containing aggregates in (**a**). Data shown are mean ± SEM. **p < 0.01, ***p < 0.001 from two-tailed, unpaired t-test or Mann Whitney test using GraphPad Prism. (**c**, **d**) WT and *NDUFA11* KO HEK293T cells were transfected with *EEF1A2*, *HSF1* or control siRNAs for 72h. (**c**) Images of protein aggregates stained with ProteoStat. Data shown are mean ± SEM. ** p < 0.01, *** p < 0.001; ns, not significant from an ordinary one-way ANOVA with Tukey’s multiple comparisons test using GraphPad Prism. (**d**) Quantification of the percentage of cells containing aggregates in (**c**). (**e**, **f**) *NDUFA11* KO and WT HEK293T cells were transfected with *PSMB9*, *HSPB1*, both *PSMB9* and *HSPB1* or control siRNAs for 72 h together with EGFP-*MAPT*. (**e**) Images of EGFP-MAPT expression. The scale bar represents 10 μm. (**f**) Quantification of the percentage of cells containing strong signals of MAPT in (**e**). (**g**) Images of PLA performed between SDHA and PSMB9 in WT and mitochondrial complex I-deficient cells. (**h**) Quantification of PLA signals per cell in (**g**). Data shown are mean ± SEM. *p < 0.05, *** p < 0.001; ND, not detected from an ordinary one-way ANOVA with Tukey’s multiple comparisons test using GraphPad Prism. The scale bar represents 10 μm. Data shown are representative of 3 independent experiments.

Some components of ubiquitin-proteasome system are localized to mitochondria (Bragoszewski, Turek et al., 2017) and CCCP treatment promotes proteasome recruitment to mitochondria (Chan, Salazar et al., 2011). Therefore, we examined whether PSMB9-containing proteasomes are recruited to mitochondria upon mitochondrial dysfunction using proximity ligation assay (PLA), a method to identify physical closeness of two proteins where a signal will only be produced if they are closer than 40 nm. For this assay, PSMB9 and a mitochondrial complex II subunit SDHA were stained to detect PSMB9-containing proteasomes and mitochondria, respectively. PLA analysis revealed that more PSMB9-containing proteasomes are closely localized to mitochondria in *NDUFA11* KO and *NDUFA13* KO than in WT HEK293T cells (Fig. 6g, h). The PSMB9-replaceable proteasome subunit PSMB6 was also stained with SDHA for PLA analysis, however, no PLA signal was detected in both mitochondrial complex I-deficient and WT HEK293T cells (Fig. 6g, h).

In sum, EEF1A2 upregulated by mitochondrial stress induces PSMB9 expression, and the induced PSMB9-containing proteasomes provide an important defense against proteotoxicity close to mitochondria in concert with HSPB1.

## DISCUSSION

Prosteostasis is challenged by mitochondrial dysfunction, and appropriate stress responses are required to sustain proteome balance. In this study, we found that HSPB1 and PSMB9 are responsible for the maintenance of proteostasis disrupted by mitochondrial dysfunction. PSMB9 induction was EEF1A2-dependent, and the induced PSMB9-containing proteasomes were localized near mitochondria, suggesting their distinct role in protein degradation associated with mitochondrial dysfunction.

Our results revealed upregulation of HSPB1 and HSPH1 mRNA and protein expression upon mitochondrial dysfunction. HSPB1 acts as a holdase by sequestering unfolded and misfolded proteins to protect them from uncontrolled aggregation in an ATP-independent manner (Haslbeck, Weinkauf et al., 2019, Mogk, Bukau et al., 2018, Mogk, Ruger-Herreros et al., 2019). HSPH1 is a substitute for HSP70 family proteins and also acts as a holdase preventing protein aggregation under stress conditions with low levels of ATP (Mogk et al., 2018, Yamagishi et al., 2003). Moreover, we found significantly more abundant HSPB1 together with 20S proteasome subunits in aggregates and purified proteasome fractions of *NDUFA11* KO compared to those of WT HEK293T cells. In *C. elegans*, enriched small HSPs and proteasome subunits were observed in insoluble fractions of long-lived mutant (Walther et al., 2015), indicating that age-related stress responses could be associated with a decline of mitochondrial function. Collectively, our study suggests that HSPB1 is required not only to prevent protein aggregation, but also required to facilitate the transfer of aggregation-prone proteins to the proteasome for degradation under mitochondrial stress conditions.

Consistent with previous findings in yeast and C.elegans (Sladowska et al., 2021, Wrobel et al., 2015), our study showed that mitochondrial deficiency enhances proteasome activity in human cells. Besides, assembly of the 20S proteasomes was promoted, and an immunoproteasome subunit PSMB9 was specifically induced among proteasome subunits upon mitochondrial dysfunction. The 20S proteasome is ATP-independent and it preferentially binds to the hydrophobic regions exposed from oxidized and damaged proteins and degrade them (Davies, 2001, Pickering & Davies, 2012, Raynes, Pomatto et al., 2016), therefore, higher levels of the 20S proteasome could be beneficial to cope with proteotoxic stress. The main purpose of the immunoproteasome was initially proposed to generate peptides for major histocompatibility complex class I antigen presentation(Murata, Takahama et al., 2018, Schmidt & Finley, 2014, Tanaka, 2009), however, growing evidence shows that the immunoproteasome preferentially degrades oxidized proteins with an activity and selectivity equal to, or greater than, those of the 20S proteasome (Raynes et al., 2016, Seifert, Bialy et al., 2010). In addition, structural differences of the constitutive proteasome and the immunoproteasome were discovered helping us to better understand different cleavage specificities of β subunits. For instance, PSMB9 preferentially cleaves after hydrophobic residues, while PSMB6 preferentially cleaves after acidic residues (Huber, Basler et al., 2012). It has been reported in recent years that mitochondrial DNA and their double-stranded RNA released from mitochondria cause immune responses such as antiviral immune responses in the cytosol (Dhir, Dhir et al., 2018, West, Khoury-Hanold et al., 2015). Based on the findings of those previous studies, we speculate that cells recognize mitochondrial proteins accumulated in the cytosol as danger signals and activate immune-like responses to defend against proteotoxicity provoked by mitochondrial stress.

In mammals, EEF1A1 is ubiquitously expressed, while EEF1A2 is normally expressed only in heart, muscle and brain (Knudsen et al., 1993, Lee, Francoeur et al., 1992) but EEF1A2 occasionally appears in some cancers (Anand, Murthy et al., 2002, Scaggiante, Dapas et al., 2012). The two isoforms have similar activities in an *in vitro* translation system, however, EEF1A2 has a greater affinity for GDP than GTP, whereas EEF1A1 has a greater affinity for GTP (Kahns, Lund et al., 1998). Despite their high amino acid sequence similarities, they exhibit differential post-translational modification, suggesting that they can be differentially regulated depending on the environmental conditions (Soares & Abbott, 2013). Beyond their major role as a translation elongation factor, they are also known to be involved in various cellular mechanisms such as cytoskeletal remodeling and signaling pathways, however, these functions might not be fully independent from their role in translation (Panasyuk, Nemazanyy et al., 2008, Soares & Abbott, 2013). It was reported that EEF1A1 participates in HSP70 expression from transcription through translation upon heat stress (Vera, Pani et al., 2014). Herein, we identified EEF1A2 as a key regulator of PSMB9 expression upon mitochondrial dysfunction.

In conclusion, we demonstrate that stress responses upregulating HSPB1 and enhancing 20S proteasome activity through PSMB9 prevent protein aggregation with minimal energetic cost under ATP-depleting conditions such as mitochondrial dysfunction. More importantly, Our data suggest that EEF1A2 participates in the selective protein expression of PSMB9 according to cellular demands, and the PSMB9-containing proteasome is located near mitochondria, enabling cells to react quickly and efficiently to degrade abnormal mitochondrial and non-mitochondrial proteins upon mitochondrial dysfunction to restore proteostasis.

## MATERIALS AND METHODS

### Cell lines and culture conditions

HEK293T wild-type and two complex I-deficient cell lines *NDUFA11*-2 KO and *NDUFA13*-2 KO, reported in Stroud et al (2016), were kindly gifted from Dr. Michael T. Ryan from Monash Biomedicine Discovery Institute, Monash University, 3800, Melbourne, Australia. The cells were cultured in Dulbecco’s modified Eagle’s medium (DMEM) with high-glucose content (4,500 mg/L) supplemented with 10% (v/v) fetal bovine serum (FBS), 2 mM L-glutamine, 1% (v/v) penicillin-streptomycin and 50 µg/ml uridine at 37°C in a 5% CO_2_ incubator. The medium was changed every other day for the maintenance of cell lines in culture and 1 day before harvest. To assess ubiquitin accumulation, cells were treated with 500nM MG132 (Enzo Life Sciences, cat. no. BML-PI102-0005) for 24h. To induce mitochondrial stress, cells were treated with 100 nM rotenone (Sigma, cat. no. R8875), 10uM menadione (Sigma, cat. no. M5625) and 10 uM CCCP (Sigma, cat. no. C2759) for 2 and 24 h. Cells added with equal volumes of DMSO were set up as control. Immortalized skin fibroblasts from a patient (mt8987i) with a missense homozygous c.221G&gt;A mutation in exon 2 of the *COX6B1* gene that causes substitution of highly conserved arginine with histidine at position 19 of the mature protein (R19H), and a healthy donor were a kind gift from Prof. Massimo Zeviani laboratory, Mitochondrial Biology Unit, MRC, Cambridge. All the fibroblasts were received with acceptance of BioBank of Telethon Italy located at the Fondazione Istituto Neurologico Carlo Besta, Milan, Italy (original source bank of cells). Immortalized skin fibroblasts were cultured at 37°C with 5% CO 2 in DMEM with high-glucose content (4,500 mg/L) supplemented with 10% (v/v) fetal bovine serum (FBS), 2 mM L-glutamine, 1% (v/v) penicillin-streptomycin, 1 mM sodium pyruvate and 50 µg/ml uridine. For the experiment, cells were seeded on 100 mm plates at density of 27,000/cm^2^ in DMEM that contained low glucose (1.1 g/l) for 24 h and after this time the medium was changed for DMEM that contained galactose (1.8 g/l) and cells were grown in this condition for the next 48 h.

### Quantitative real-time PCR

Total RNA was isolated with RNeasy Plus Mini Kit (Qiagen, cat. no. 74134) according to the manufacturer’s instructions. 1 ug total RNA was used to generate cDNA using SuperScript™ IV First-Strand Synthesis System (Thermo Fisher Scientific, cat. no. 18091050). To measure mRNA stability, cells were treated with 5 µg/ml Actinomycin D (Sigma, cat. no. A9415) to inhibit transcription. Cells added with equal volumes of DMSO were set up as control. After 0, 4, 8, 16 and 24 h of Actinomycin D treatment, 1 ug total RNA was extracted and cDNA was synthesized using Maxima First Strand cDNA Synthesis Kit for RT-qPCR, with dsDNase (Thermo Fisher Scientific, cat. no. K1671). RT-qPCR was performed using SensiFAST™ SYBR® Hi-ROX Kit (Bioline, cat. no. BIO-92020) in a 96-well white plate (Roche, cat. no. 4729692001) using a LightCycler480 (Roche) with two technical repeats per experimental condition. Fold changes in mRNA expression of the target genes were calculated using the ΔΔCt method. The expression levels of ACTB were used as internal controls.

### RNA-seq sample preperation

Total RNA was isolated with RNeasy Plus Mini Kit (Qiagen, cat. no. 74134) followed by DNase I (Sigma, cat. no. AMPD1) treatment. Twelve total RNA samples were subjected to mRNA enrichment using mRNA NEBNext® Poly(A) mRNA Magnetic Isolation Module (New England Biolabs, cat. no. E7490L), from which libraries were derived. Libraries were constructed with KAPA RNA HyperPrep Kit (Kapa Biosciences, cat. no. 08098107702) and KAPA Dual-Indexed adapter kit (Kapa Biosciences, cat. no. KK8722), following manufacturer’s standard protocols, with 2μg RNA as input, 5 min fragmentation in 94°C and 11 cycles of amplification. The quality of obtained libraries were analyzed using Agilent Bioanalyzer 2100 and High Sensitivity DNA kit (Agilent, cat. no. 5067-4626). Finally, the quantity was measured by qPCR with Kapa Library Quantification kit (Kapa Biosciences, cat. no. KK4824), according to the manufacturer’s instructions. Sequencing was performed on Illumina NovaSeq 6000 using NovaSeq 6000 S1 Reagent Kit (200 cycles) (Illumina, cat. no. 20012864), with pair-end mode of 2×100 cycles and standard operating procedure.

### RNA-seq analysis

Raw sequences were trimmed according to quality using Trimmomatic(Bolger, Lohse et al., 2014) (version 0.39) using default parameters, except MINLEN, which was set to 50. Trimmed sequences were mapped to human reference genome provided by ENSEMBL, (version grch38_snp_tran) using Hisat2 (Kim, Langmead et al., 2015) with default parameters. Optical duplicates were removed using MarkDuplicates tool from GATK(McKenna, Hanna et al., 2010) (version 4.1.2.0) with default parameters except OPTICAL_DUPLICATE_PIXEL_DISTANCE set to 12000. Reads that failed to map to the reference were extracted using Samtools (Li, Handsaker et al., 2009) and mapped to Silva meta-database of rRNA sequences (Quast, Pruesse et al., 2013) (version 119) with Sortmerna(Kopylova, Noe et al., 2012) (version 2.1b) using “–best 1” option. Mapped reads were associated with transcripts from GRCh38 database (Zerbino, Achuthan et al., 2018) (Ensembl, version 77) using HTSeq-count (Anders, Pyl et al., 2015) (version 0.9.1) with default parameters except –stranded set to “reverse”. Differentially expressed genes were selected using DESeq2 package(Love, Huber et al., 2014) (version 1.16.1). Fold change was corrected using apeglm (Zhu, Ibrahim et al., 2019). Full lists of genes for *NDUFA11* KO and *NDUFA13* KO samples versus WT samples are available in Supplementary Figure 1-Source Data 1. Only genes with a maximum Benjamini-Hochberg corrected *p* value (q-value) of 0.05 with at least combined mean of 32 reads were deemed to be significantly differentially expressed.

### Reactome pathway enrichment analysis

Reactome pathways enrichment analysis was performed using R Biocunductor package ReactomePA version 1.36 (Yu & He, 2016). The calculations were done with enrichedPathway() function using default argument values - minimal size of genes annotated by Ontology term for testing was set to 10, p-value cutoff was set to 0.05 and q-value cutoff for 0.2. p values were adjusted for multiple testing using FDR controlling method of Benjamini and Hochberg (Benjamini & Hochberg, 1995).

### siRNA-mediated knockdown

To silence endogenous *PSMB6*, *PSMB9* or *HSPB1* expression, ON-TARGETplus SMARTpool siRNAs targeting PSMB6 (Dharmacon, cat. no. L-006020-00-0020), *PSMB9* (Dharmacon, cat. no. L-006023-00-0050), *HSPB1* (Dharmacon, cat.no. L-005269-00-0050) were used; ON-TARGETplus Non-targeting siRNA (Dharmacon, cat. no. D-001810-10-50) was used as a negative control. To silence endogenous *EEF1A1*, *EEF1A2* or *HSF1* expression, Silencer™ Pre-Designed siRNAs targeting EEF1A1 (Ambion, ID: 2991), EEF1A2 (Ambion, ID: 10789) or *HSF1* (sense strand: 5’-CGGAUUCAGGGAAGCAGCUGGUGCA-3’, Sigma) were used; Silencer™ Negative Control No. 1 siRNA (Ambion, cat. no.AM4635) was used as a negative control. Cells were reverse-transfected with 20nM of siRNA using Lipofectamine™ RNAiMAX Transfection Reagent (Thermo Fisher Scientific, cat. no. 13778150) diluted in Opti-MEM™ I Reduced Serum Medium (Thermo Fisher Scientific, cat. no. 11058021) according to the manufacturer’s instructions. Three days after transfection, cells were collected for subsequent analysis.

### Plasmids and transfection

To generate an expression vector for FLAG-tagged *PSMA5*, cDNA fragment containing the entire protein encoding ORF of PSMA5 was amplified with a forward primer containing a EcoRI site (5’-TTTT GAATTC ATG TTT CTT ACC CGG TCT GAG TAC GAC-3’, Sigma) and a reverse primer containing sequences specifying an FLAG tag, a stop codon, and an EcoRI site (5′-TTTT GAATTC TTA CTT ATC GTC GTC ATC CTT GTA ATC AAT GTC CTT GAT AAC CTC TTC AAG-3’, Sigma). The PCR products were digested with EcoRI restriction enzyme and cloned into the EcoRI-digested pcDNA3.1/Zeo(+) vector. Cloned construct was sequenced for insert verification. Cells were transfected with GeneJuice® Transfection Reagent (Merck Millipore, cat. no.70967) according to the manufacturer’s instructions. Control cells were transfected with empty pcDNA3.1/Zeo(+). Three days after transfection, cells were collected for subsequent analysis. Expression plasmid containing Myc-DDK-tagged *NDUFA11* cDNA was purchased from Origene (Origene, cat. no. RC208966) and pRK5-EGFP-*MAPT* was a gift from Karen Ashe (Addgene plasmid # 46904 ; http://n2t.net/addgene:46904 ; RRID:Addgene_46904). Control cells were treated with transfection reagents. Double concentration of plasmid was applied to *NDUFA11* KO for MAPT expression to equalize protein expression in WT HEK293T cells due to its low translation efficiency. Three days after transfection, cells were collected for subsequent analysis.

### Western blot analysis

Cells were lysed in RIPA buffer (65 mM Tris base pH 7.4, 150 mM NaCl, 1% (v/v) NP-40, 0.25% sodium deoxycholate, 1 mM EDTA and 2 mM phenylmethylsulfonyl fluoride (PMSF)) for 30min at 4°C. The lysate was clarified by centrifugation at 14,000 x g for 30 min at 4°C. The supernatant was collected and the protein concentration was measured by the Bradford protein assay.The supernatant was diluted in Laemmli buffer that contained 50mM dithiothreitol and denatured at 65°C for 15min. Total protein extracts were resolved on 8% or 15% SDS-PAGE gels, transferred to PVDF membranes (Sigma, cat. no. GE10600021) and probed with specific primary and secondary antibodies, and developed by standard techniques using chemiluminescence. The primary antibodies used for immunoblotting were as follows: HSPB1 (Abcam, cat. no. ab2790, 1:500), HSPA1A/HSP1AB (Enzo Life Sciences, cat. no. ADI-SPA-812-F, 1:2,000), HSP90 (Abcam, cat. no. ab13495, 1:1,000), HSPH1 (Abcam, cat. no. ab109624, 1:500), ACTB (Sigma, cat. no. A1978, 1:2,000), NDUFA11 (Abcam, cat. no. ab183707, 1:500), NDUFA13 (Abcam, cat. no. ab110240), PSMB5 (Enzo Life Sciences, cat. no. BML-PW8895-0100, 1:500), PSMB6 (Abcam, cat. no. ab150392, 1:500), PSMB8 (Abcam, cat. no. ab3329, 1:500), PSMB9 (Abcam, cat. no. ab150392, 1:500), PSMD1 (Abcam, cat. no. ab2941, 1:4,000), Proteasome 20S alpha 1+2+3+5+6+7 antibody (Abcam, cat. no. ab22674, 1:1,000), Ubiquitin (Santa cruz, cat. no. sc-8017), EEF1A1 (Proteintech, cat. no. 11402-1-AP, 1:1,000), EEF1A2 (Thermo Fisher Scientific, cat. no. PA5-27677, 1:500), FLAG (Sigma, cat. no. F1804, 1:500), MAPT (Sigma, cat. no. 577801, 1:2,000). The secondary horseradish peroxidase-conjugated anti-mouse (Sigma, cat. no. A4416, 1:5,000 or 1:10,000) and anti-rabbit (Sigma, cat. no. A6154, 1:5,000 or 1:10,000) were used.

### Proteasome in-solution activity assay and native gel electrophoresis of proteasome complexes

Cells were lysed in proteasome lysis buffer (50 mM Tris–HCl, pH 7.4, 10 mM MgCl_2_, 250 mM sucrose, 0.5 mM EDTA, 2 mM ATP, 1 mM DTT and 2 mM PMSF) and homogenized in a dounce glass homogenizer. The homogenate was clarified by centrifugation at 10,000 x g for 15 min at 4°C, and protein concentration was quantified using the Bradford protein assay. 4 ug protein were incubated with 50 uM Suc–Leu–Leu–Val–Tyr– AMC peptide substrate (chymotrypsin-like activity; Bachem, cat. no. I-1395) or 50 uM Ac– NIe–Pro–NIe–Asp–AMC (caspase-like activity; Bachem, cat. no. I-1850) and 10ug protein were incubated with 50 uM Ac-Pro-Ala-Leu-AMC (Cayman Chemical, cat. no. 26592) to measure the activity of PSMB9 in a final volume of 200 ml of lysis buffer in a 96-well plate. Fluorescence (excitation wavelength 380 nm, emission wavelength 460 nm) was measured every 5 min for 2 h at 25°C using Synergy H1 Hybrid Multi-Mode Microplate Reader (BioTek, cat. no. H1MFDG). The rate of kinetic reaction (slope) of proteasome activity was calculated and data are represented in a fold change compared to wild-type cells. For native gel electrophoresis, 50ug of proteins were separated on 4.5% native polyacrylamide gel that was supplemented with 5mM MgCl_2_, 2.5% sucrose and 1 mM ATP. Electrophoresis was performed at 4°C for 3.5 h at 30 min in running buffer (90 mM Tris base, 90 mM boric acid, 0.5 mM EDTA pH 8.0, 5 mM MgCl_2_ and 1 mM ATP).

### Immunopurification of proteasomes

Cells transfected with FLAG-tagged PSMA5 expression plasmid or empty vector were lysed in proteasome lysis buffer and protein concentration was determined as described above. 8 mg protein were incubated with ANTI-FLAG® M2 Affinity Gel (Sigma, cat.no. A2220) for 2 h on a rotator at 4°C. Immunoprecipitates were washed three times with wash buffer (20 mM Tris-HCl ph7.4, 150 mM NaCl, 1 mM EDTA and 2 mM PMSF) and eluted with 1 mg/ml FLAG® Peptide (Sigma, cat. no. F3290) in wash buffer overnight on a rotator at 4°C for anaylsis by mass spectrometry.

### IP after MAPT expression

Cells were transfected with EGFP-tagged MAPT expression plasmid for 72 h, harvested in ice-cold PBS, centrifuged at 1,000 x g for 3 min at 4°C, and washed twice with ice-cold PBS. Cell pellet was resuspended in 500 μl of ice-cold lysis buffer (10 mM Tris/Cl pH 7.5, 150 mM NaCl, 0.5 mM EDTA, 0.5 % IGEPAL CA-630 (NP-40), and 2 mM PMSF) by pipetting and incubated on ice for 30 min with extensively pipetting every 10 min. The cell lysate was centrifuged at 20 000 x g for 10 min at 4°C and then the supernatant was diluted with wash buffer (10 mM Tris-HCl pH 7.5, 150 mM NaCl and 0.5 mM EDTA) to obtain 850 µg of protein in 500 µl of the lysate. GFP-Trap®_A beads (25 µl) (Chromotek, cat. no. gta-20) were centrifuged at 2500 x g for 2 min at 4°C and washed twice with ice-cold wash buffer. Then, 500 µl of diluted cell lysate was added to prepared GFP-Trap®_A beads and incubated with rotation overnight at 4°C. Next, GFP-Trap®_A beads were centrifuged at 2500 x g for 2 min at 4°C and washed 6 times with ice-cold wash buffer. GFP-Trap®_A beads were finally resuspended in 50 μl 2X SDS-Sample buffer and boiled for 10 min at 95°C to dissociate the immunocomplexes from the beads. The beads were centrifuged at 2,500 x g for 2 min at 4°C and collected supernatant was separated by SDS–PAGE on 15 % gel.

### Preparation of aggregate samples for proteomic analysis

Isolated protein aggregate fraction was solubilized for 1 h at room temperature with the use of 10 µl 0.1 % ProteaseMAX™ surfactant (Promega, cat. no. V2072) in 50 mM ammonium bicarbonate. Then, 50 µl of digestion mixture (50 mM Tris-Cl pH 8.2; 5 mM TCEP; 10 mM iodoacetamide; 0.5 µg of sequencing grade modified trypsin (Promega, cat.no. V5111) and 0.02 % ProteaseMAX™ surfactant) was added and the sample was digested overnight at 37°C with shaking (850 rpm) in low protein binding tubes. Next, the sample was diluted with 100 µl of H 2 O 2, and trifluoroacetic acid (TFA) was added to a final concentration of 1 % in order to inactivate the trypsin. Then, the sample was mixed in the thermomixer at 850 rpm for 2 minutes at 37°C and centrifuged at 12,000 x g for 3 min at room temperature. The supernatants were then desalted with the use of AttractSPE™ Disks Bio – C18 (Affinisep, cat. no. SPE-Disks-Bio-C18-100.T1.47.20) using a published stage-tip protocol (Rappsilber, Mann et al., 2007), and concentrated using a Savant SpeedVac concentrator. Prior to LC-MS measurement, the samples were resuspended in 0.1 % TFA, 2% acetonitrile in water.

### LC-MS/MS analysis for protein aggregates

Chromatographic separation was performed on an Easy-Spray Acclaim PepMap column 15cm long × 75 µm inner diameter (Thermo Fisher Scientific) at 35 °C by applying a 90 min acetonitrile gradients in 0.1% aqueous formic acid at a flow rate of 300 nl/min. An UltiMate 3000 nano-LC system was coupled to a Q Exactive HF-X mass spectrometer via an easy-spray source (all Thermo Fisher Scientific). The Q Exactive HF-X was operated in data-dependent mode with survey scans acquired at a resolution of 60,000 at m/z 200. Up to 12 of the most abundant isotope patterns with charges 2-6 from the survey scan were selected with an isolation window of 1.3 m/z and fragmented by higher-energy collision dissociation (HCD) with normalized collision energies of 27, while the dynamic exclusion was set to 30 s. The maximum ion injection times for the survey scan and the MS/MS scans (acquired with a resolution of 15,000 at m/z 200) were 45 and 22 ms, respectively. The ion target value for MS was set to 3e6 and for MS/MS to 10e5, and the minimum AGC target was set to 8e2.

### Proteomics data processing for protein aggregates

The data were processed with MaxQuant v. 1.6.7.0 (Cox & Mann, 2008), and the peptides were identified from the MS/MS spectra searched against Uniprot KB Human Proteome using the build-in Andromeda search engine. Raw files obtained from the LC-MS/MS measurements of protein aggregates originated from *NDUFA11* KO and WT HEK293T cells in the absence or presence of EGFP-MAPT were processed together (4 groups, 3 samples each). Cysteine carbamidomethylation was set as a fixed modification and methionine oxidation as well as protein N-terminal acetylation were set as variable modifications. For in silico digests of the reference proteome, cleavages of arginine or lysine followed by any amino acid were allowed (trypsin/P), and up to two missed cleavages were allowed. The FDR was set to 0.01 for peptides, proteins and sites. Match between runs was enabled. Other parameters were used as pre-set in the software. Unique and razor peptides were used for quantification enabling protein grouping (razor peptides are the peptides uniquely assigned to protein groups and not to individual proteins). Data were further analysed using Perseus version 1.6.6.0 and Microsoft Office Excel 2016.

### LFQ-based differential analysis of aggregated protein level

LFQ values for protein groups were loaded into Perseus v. 1.6.6.0. Standard filtering steps were applied to clean up the dataset: reverse (matched to decoy database), only identified by site, and potential contaminant (from a list of commonly occurring contaminants included in MaxQuant) protein groups were removed. LFQ intensities were log2 transformed and protein groups with log2 LFQ values for at least 2 samples in at least one experimental group were kept. Further analysis was performed in 2 experimental group pairs: 1) *NDUFA11* KO vs WT HEK293T cells in the absence of EGFP-MAPT; 2) *NDUFA11* KO vs WT HEK293T cells in the presence of EGFP-MAPT. Within each experimental group pair, protein groups with log2 LFQ values for at least 2 samples in both experimental groups were kept, while protein groups with log2 LFQ values for at least 2 samples in one experimental group and log2 LFQ value for exactly 1 sample in the other group were discarded. For protein groups with log2 LFQ values for at least 2 samples in one experimental group and no log2 LFQ values in the other, data imputation from normal distribution (width = 0.3, down shift = 1.8 × standard deviation) was performed within the experimental group with 0 values. Gaussian distribution of log2 LFQ intensities were confirmed by histogram analysis preventing the unbiased introduction of small values. Student’s T-test (permutation-based FDR = 0.05, S0 = 0.1) was performed to return proteins which levels were statistically significantly changed in response to *NDUFA11* KO.

### Quantitative Mass Spectrometry for proteasome complexes

Proteins of proteasome complexes affinity-purified via FLAG-tagged PSMA5 from *NDUFA11* KO, *NDUFA13* KO or WT (control) cells were acetone-precipitated and resuspended in 6 M urea/50 mM ammonium bicarbonate, followed by reduction and alkylation of cysteine residues and tryptic in solution digestion as described before (Peikert, Mani et al., 2017). Peptides were differentially labeled by stable isotope dimethyl labeling as described previously(Dannenmaier, Stiller et al., 2018) using ‘light’ formaldehyde and sodium cyanoborohydride (CH2O/NaBH3CN) for peptides originating from WT cells and the corresponding deuterated ‘heavy’ versions (CD2O/NaBD3CN) for peptides from *NDUFA11* KO and *NDUFA13 KO* cells. Labeling efficiencies were assessed by liquid chromatography-mass spectrometry (LC-MS). Equal amounts of light and heavy labeled peptides were mixed, desalted on StageTips, and analyzed by LC-MS (n = 3 per experiment, i.e. *NDUFA11* KO versus WT and *NDUFA13 KO* versus WT) using either an Orbitrap Elite or a Q Exective Plus mass spectrometer, each connected to an UltiMate 3000 RSLCnano HPLC system. Mass spectrometric raw data were searched against the Uniprot human proteome set including isoforms, to which the sequence for PSMA5-FLAG was added, and a list of common contaminants using MaxQuant/Andromeda (version 1.6.0.1; (Cox & Mann, 2008, Cox, Neuhauser et al., 2011)). MaxQuant was operated with default settings, except that protein identification and quantification were based on ≥ 1 unique peptide and ≥ 1 ratio count, respectively. DimethylLys0/dimethylNter0 and dimethylLys6/dimethylNter6 were selected as light and heavy labels, carbamidomethlyation of cysteine residues was set as fixed modification, N-terminal acetylation and oxidation of methionine were considered variable modifications, and ‘match between runs’ and ‘requantify’ were enabled. To evaluate effects of *NDUFA11* KO or *NDUFA13* KO on the composition of the proteasome, the mean log2 of normalized protein abundance ratios (i.e. PSMA5 complexes affinity-purified from *NDUFA11* KO or *NDUFA13 KO* versus WT cells) and the corresponding p-values (Student’s t-test, two-sided) were determined.

### Immunofluorescence

Cells were seeded on glass coverslips coated with 50 µg/ml poly-D-lysine (Sigma, cat. no. P7280). After 24 h, cells were fixed with 3.7% formaldehyde for 20 min and permeabilized with PBS containing 0.1% Triton X-100 for 10 min. Proximity ligation assay (PLA) was subsequently performed using Duolink® In Situ Detection Reagents Orange (Sigma, DUO92007) according to the manufacturer’s instructions using primary antibodies against PSMB9 (Abcam, cat. no. ab150392, 1:500) and SDHA (Santa Cruz Biotechnology, cat. no. sc-166947, 1:100). To visualize protein aggregates, PROTEOSTAT® Aggresome detection kit (Enzo Life Sciences, cat. no. ENZ-51035) was used following the manufacturer’s instructions. For MAPT detection, cells transfected with EGFP-MAPT expression plasmid were fixed with 3.7% formaldehyde for 20 min. After mounting of the coverslips with ProLong Diamond antifade mountant with DAPI (Thermo Fisher Scientific, P36962) on a microscope slide, images were acquired by a Zeiss LSM700 confocal microscope using 405 nm laser for DAPI, 488 nm laser for protein and MAPT aggregates, and 555 nm laser for PLA signals using a 20x objective up to 80x with a digital zoom. The total number of cells in the images taken in 8∼11 different regions at 20x magnification was automatically calculated by ImageJ, and cells with aggregates in the same images were manually counted for quantification of protein aggregates. For quantification of MATP aggregation, the total number of cells in the images taken in 5 different regions at 20x magnification was automatically calculated by ImageJ, and cells with MAPT aggregates in the same images were manually counted. To quantify PLA signals, the total number of cells in the images taken in 14 different regions for PSMB9 and SDHA staining, and 9 different regions for PSMB6 and SDHA staining at 80x magnification were manually calculated and PLA signals in the same images were automatically calculated by ImageJ.

### Isolation of protein aggregates

Cells were resuspended in lysis buffer (30 mM Tris-HCl pH 7.4, 20 mM KCl, 150 mM NaCl, 5 mM EDTA, 1% Triton X-100 and 2 mM PMSF). After incubation on ice for 30 min (with vortexing every 10 min for 10 sec), samples were shortly centrifuged at a low speed (4°C, 1000 x g, 1 min) to remove cellular debris. The supernatant was collected and the protein concentration was determined using a Bradford protein assay. 500 ug protein were further centrifuged at a high speed (4°C, 125,000 x g, 1 h) and separated in soluble and pellet fractions.

### Statistical analysis

For the statistical analysis, two-tailed, unpaired t-tests assuming equal or unequal variance were used unless stated otherwise. Values of p-value < 0.05 were considered statistically significant. Data are presented as mean values ± SD of three independent biological replicates unless stated otherwise.

## ACKNOWLEDGEMENTS

We thank David A. Stroud and Michael T. Ryan, Department of Biochemistry and Molecular Biology, Monash Biomedicine Discovery Institute, Monash University, for providing *NDUFA11* KO, *NDUFA13* KO and WT HEK293T cells, Massimo Zeviani, MRC Mitochondrial Biology Unit, University of Cambridge, for providing the immortalized fibroblasts with *COX6B1* mutation from a patient. The biobank “Cell Line and DNA Bank of Genetic Movement Disorders and Mitochondrial Diseases”, a member of the Telethon Network of Genetic Biobanks (project no. GTB12001), funded by Telethon Italy, and the EuroBioBank Network provided the control and patient-derived fibroblast specimens. We also thank Silke Oeljeklaus, Faculty of Biology, Biochemistry and Functional Proteomics, Institute of Biology II, University of Freiburg, for assistance in writing of proteomic methods. The work was funded by the ‘Regenerative Mechanisms for Health’ project MAB/2017/2, performed within the International Research Agendas program of the Foundation for Polish Science, co-financed by the European Union under the European Regional Development Fund and the National Science Centre grant 2015/18/A/NZ1/00025. NGS was performed thanks to Genomics Core Facility CeNT UW, using NovaSeq 6000 platform financed by Polish Ministry of Science and Higher Education (decision no. 6817/IA/SP/2018 of 2018-04-10).

## AUTHOR CONTRIBUTIONS

M.K. and A.C. conceived and designed the study. M.K. performed most of the experiments. M.K. and A.C. evaluated the data. L.S. performed IP analysis. L.S and M.K performed aggregation assay. T.M.S. performed the RNA-seq visualization. B.W. supervised proteomic analysis and I.S. performed proteomic analysis of purified proteasomes. R.S. performed the proteomic analysis of protein aggregates. A.K. performed western blot analysis of human fibroblasts. M.K. prepared the figures. M.K. and A.C. interpreted the results and wrote the manuscript with input from all authors.

---

**Supplementary Figure 1.**
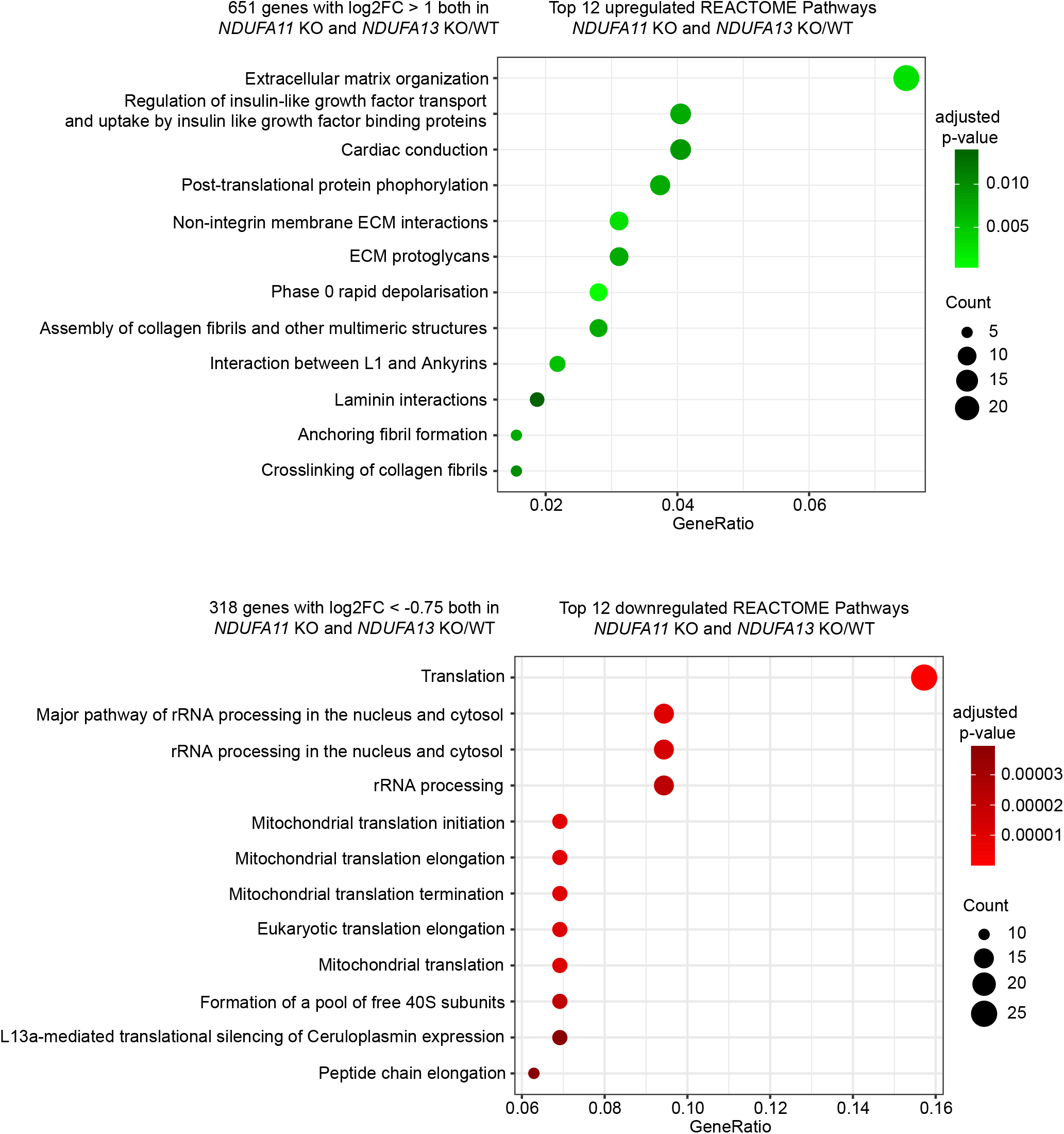
Mitochondrial complex I deficiency causes transcriptome changes. Top 12 pathways of the Reactome pathway enrichment analysis for genes upregulated with log2 fold change (log2FC) > 1 and q-value < 0.05 (upper panel, green) and downregulated with log2 fold change (log2FC) < −.75 and q-value < 0.05 (lower panel, red) in both *NDUFA11* KO and *NDUFA13* KO compared to WT HEK-293T cells.

**Supplementary Figure 2.**
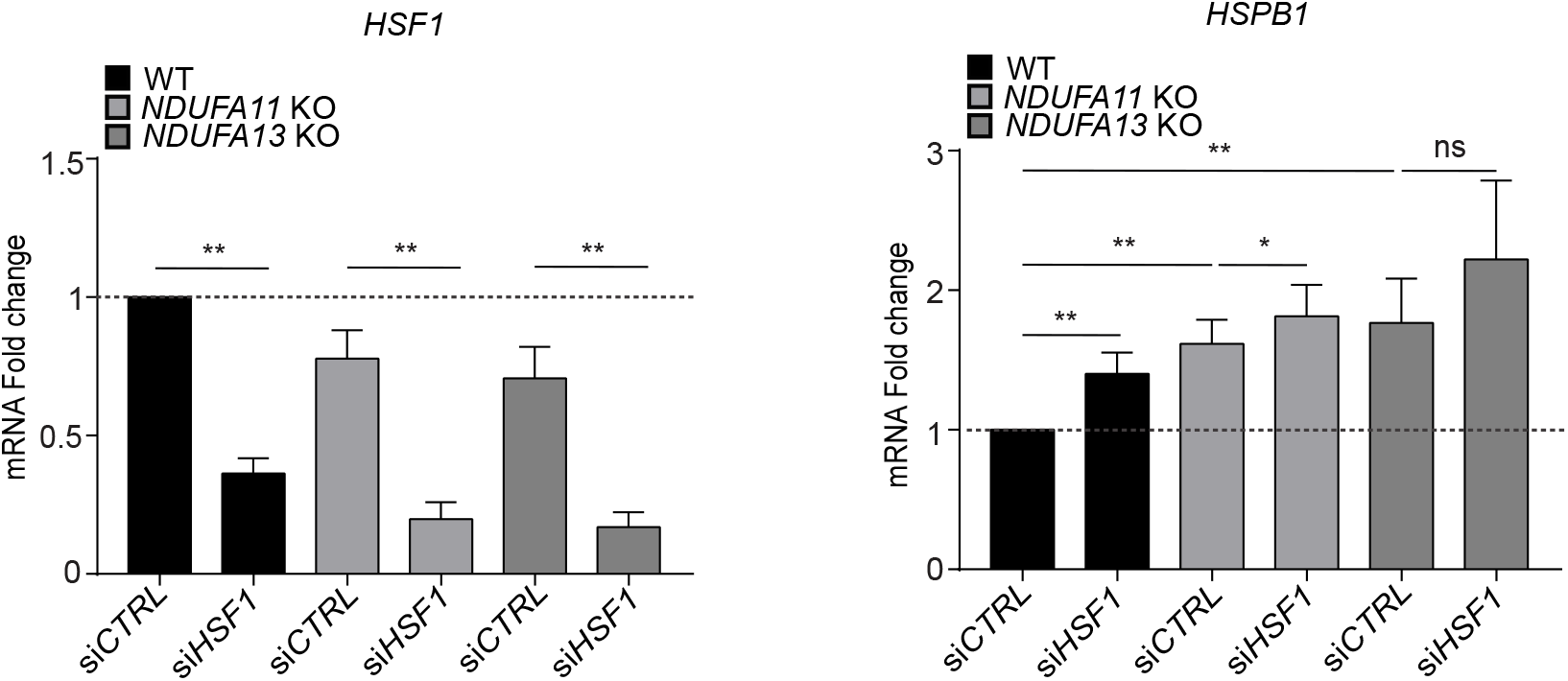
HSF1 does not regulate *HSPB1* gene expression under mitochondrial stress. mRNA expression levels of *HSF1* and *HSPB1* examined by RT-qPCR analysis in mitochondrial complex I-deficient HEK293T cells and WT HEK293T cells transfected with *HSF1* or control siRNAs for 72 h. *p < 0.05, **p < 0.01; ns, not significant from two-tailed, unpaired Mann Whitney test using GraphPad Prism.

**Supplementary Figure 3.**
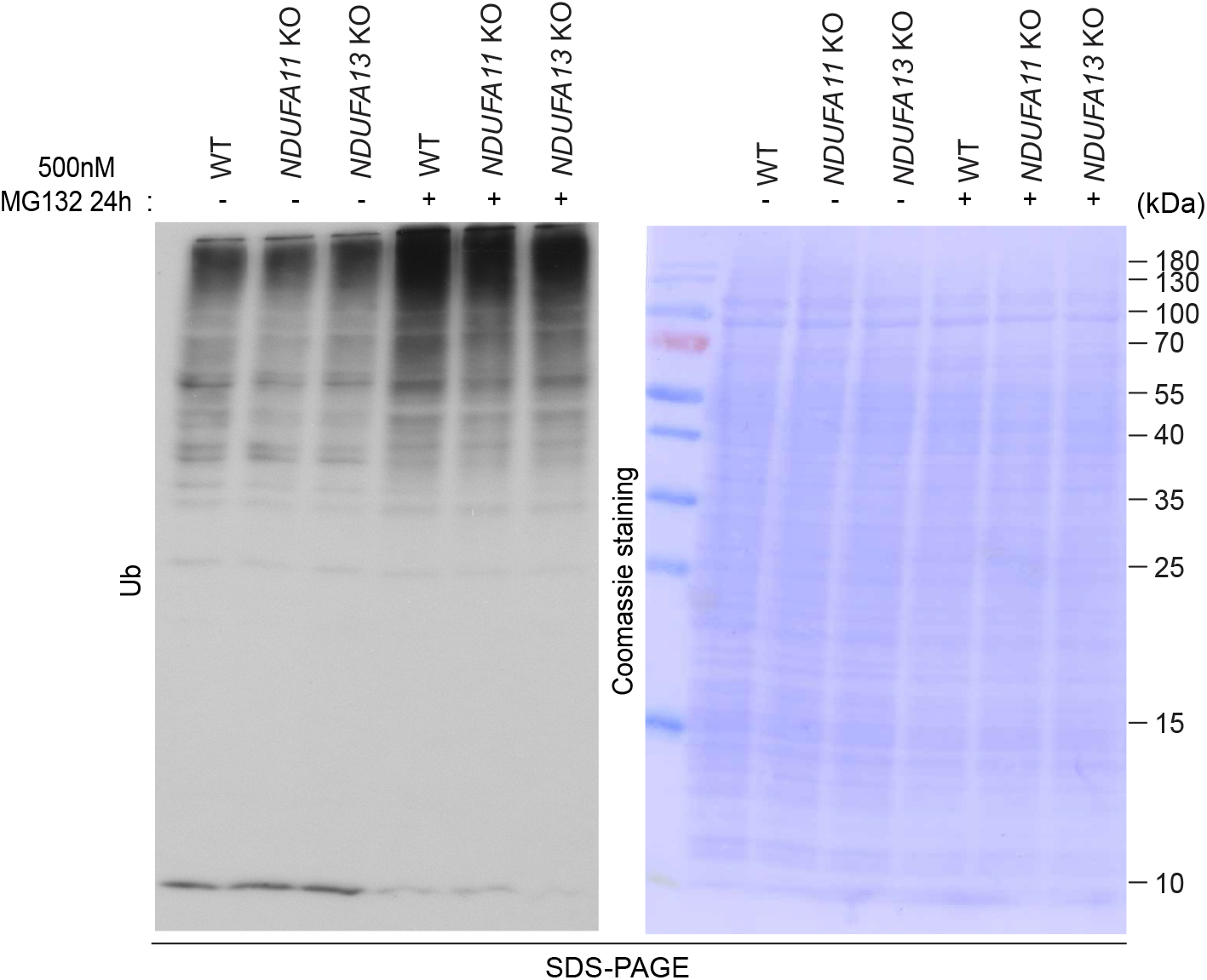
Increased proteasome activity in mitochondrial complex I-deficient cells functions ubiquitin-independently. Western blot analysis performed in whole cell lysates of mitochondrial complex I-deficient HEK293T cells and WT HEK293T cells after 24 h of MG132 treatment. Coomassie Blue staining was used as a loading control. Data shown are representative of 3 independent exper**i**ments.

**Supplementary Figure 4.**
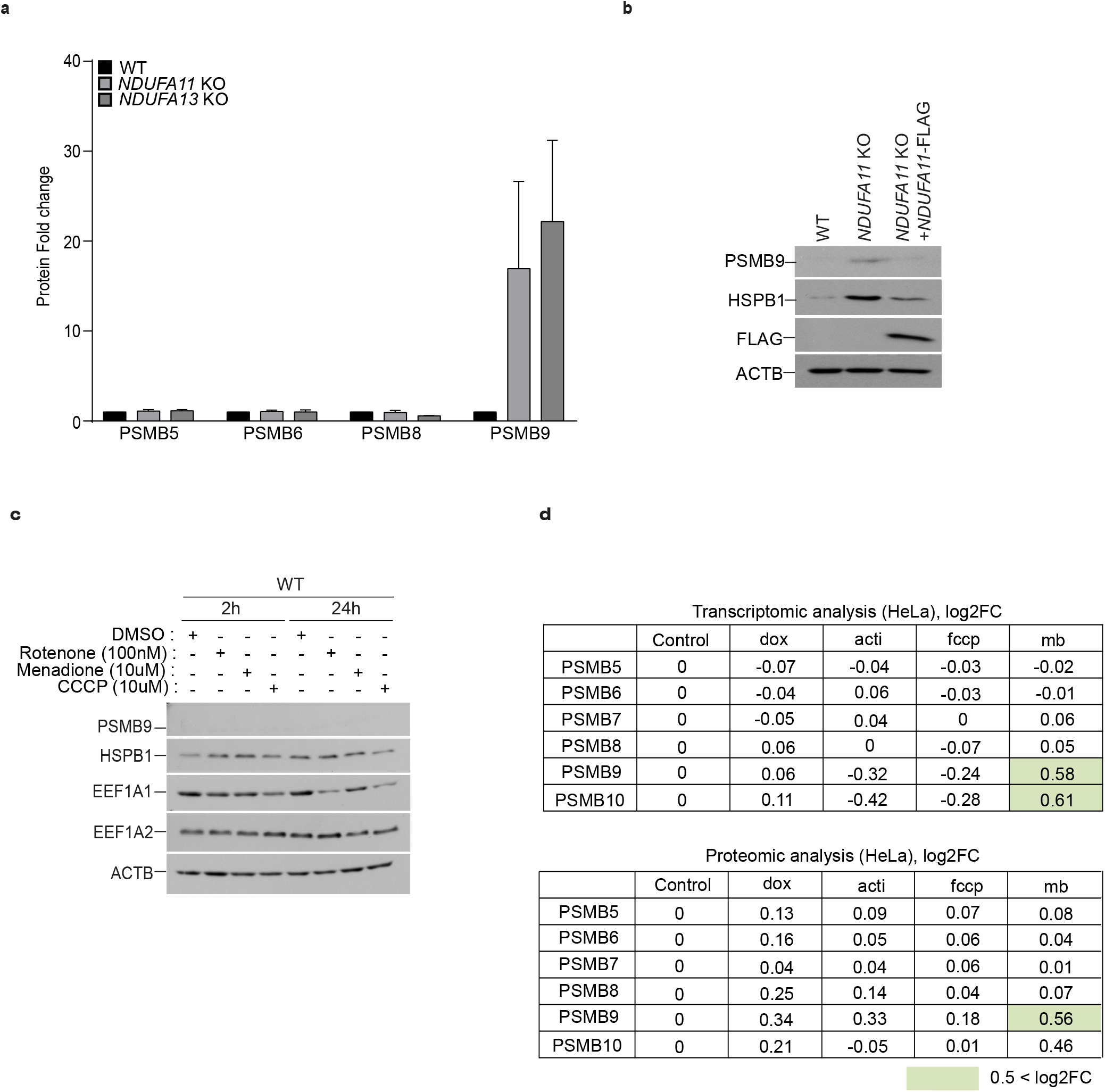
Mitochondrial import stress may trigger upregulation of immunoproteasomes. (**a**) Quantification of β subunits in western blot analysis normalized to ACTB using ImageJ. The protein levels are presented as fold changes relative to WT. Data shown are mean ± SD. (**b**) Western blot analysis performed in whole cell lysates of *NDUFA11* KO and WT HEK293T treated with transfection reagents, and *NDUFA11* KO HEK293T cells transfected with FLAG-tagged NDUFA11 expression plasmid. (**c**) Western blot analysis performed in whole cell lysates of WT treated with DMSO, rotenone, menodione or CCCP for 2 and 24 h. ACTB was used as a loading control. Data shown are representative of 3 independent experiments. (**d**) Analysis of published transcriptomic and proteomic data of HeLa cells after treatment of doxycycline (dox), actinonin (acti), fccp or MitoBlock-6 (mb) (n=2). Upregulated genes and proteins with log2 fold change (log2FC) > 0.5 are shown in green.

**Supplementary Figure 5.**
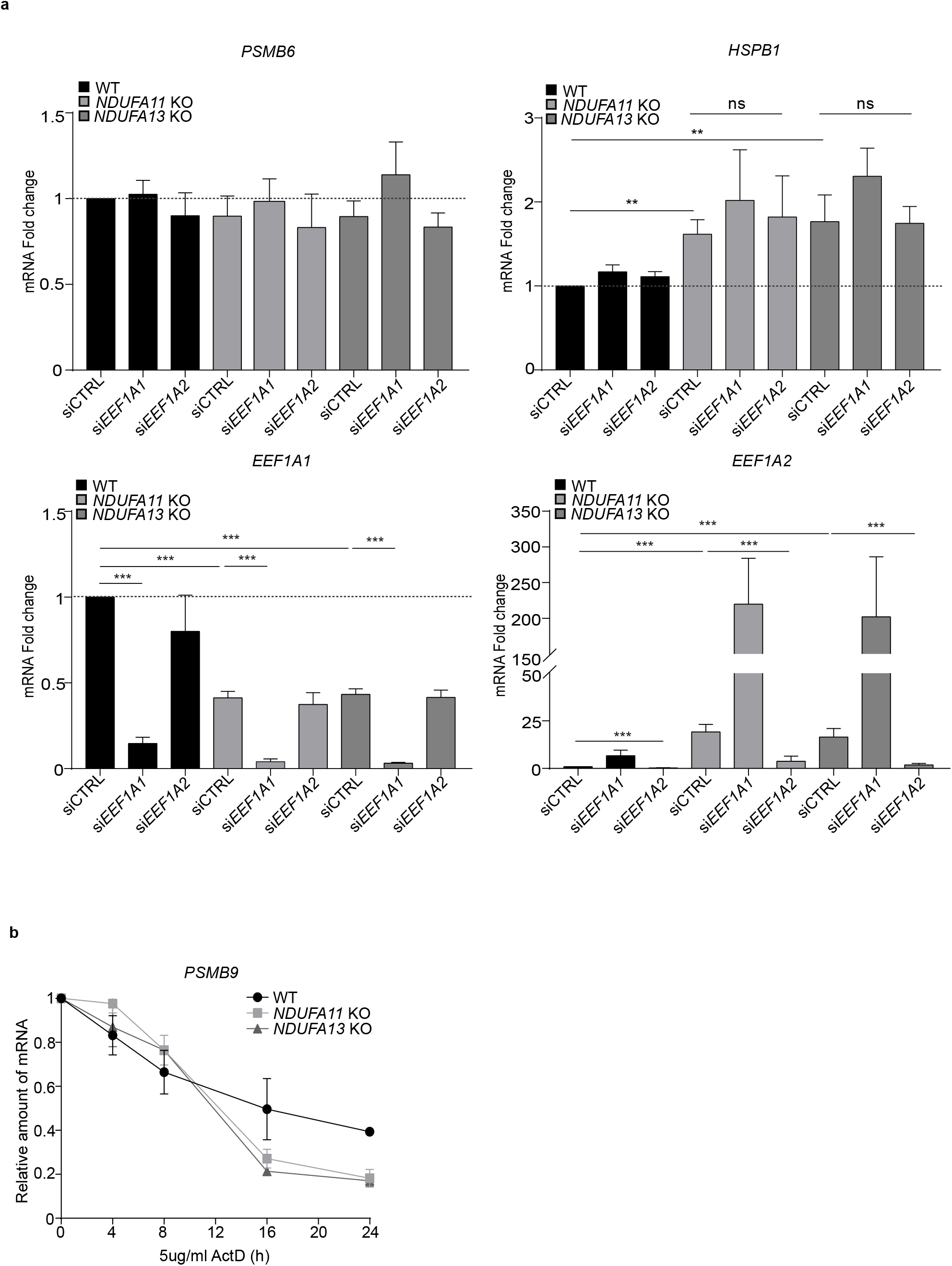
*PSMB6* and *HSPB1* mRNA expressions are EEF1A2-independent, and EEF1A2 is not involved in *PSMB9* mRNA stability. (**a**) *PSMB6*, *HSPB1*, *EEF1A1* and *EEF1A2* mRNA expression patterns examined by RT-qPCR in mitochondrial complex I-deficient and WT HEK293T cells transfected with *EEF1A1*, *EEF1A2* or control siRNAs for 72 h. The mRNA levels are presented as fold changes relative to WT transfected with control siRNA. Data shown are mean ± SD. **p < 0.01, ***p < 0.001; ns, not significant from two-tailed, unpaired t-test or Mann Whitney test using GraphPad Prism. (**b**) *PSMB9* mRNA stability assay performed by RT-qPCR after 0, 4, 8, 16 and 24h Actinomycin D treatment in *NDUFA11* KO and WT HEK293T cells. Data are presented as fold changes of mRNA relative to DMSO-treated control at each time point and are mean ± SD (n=2).

**Supplementary Figure 6.**
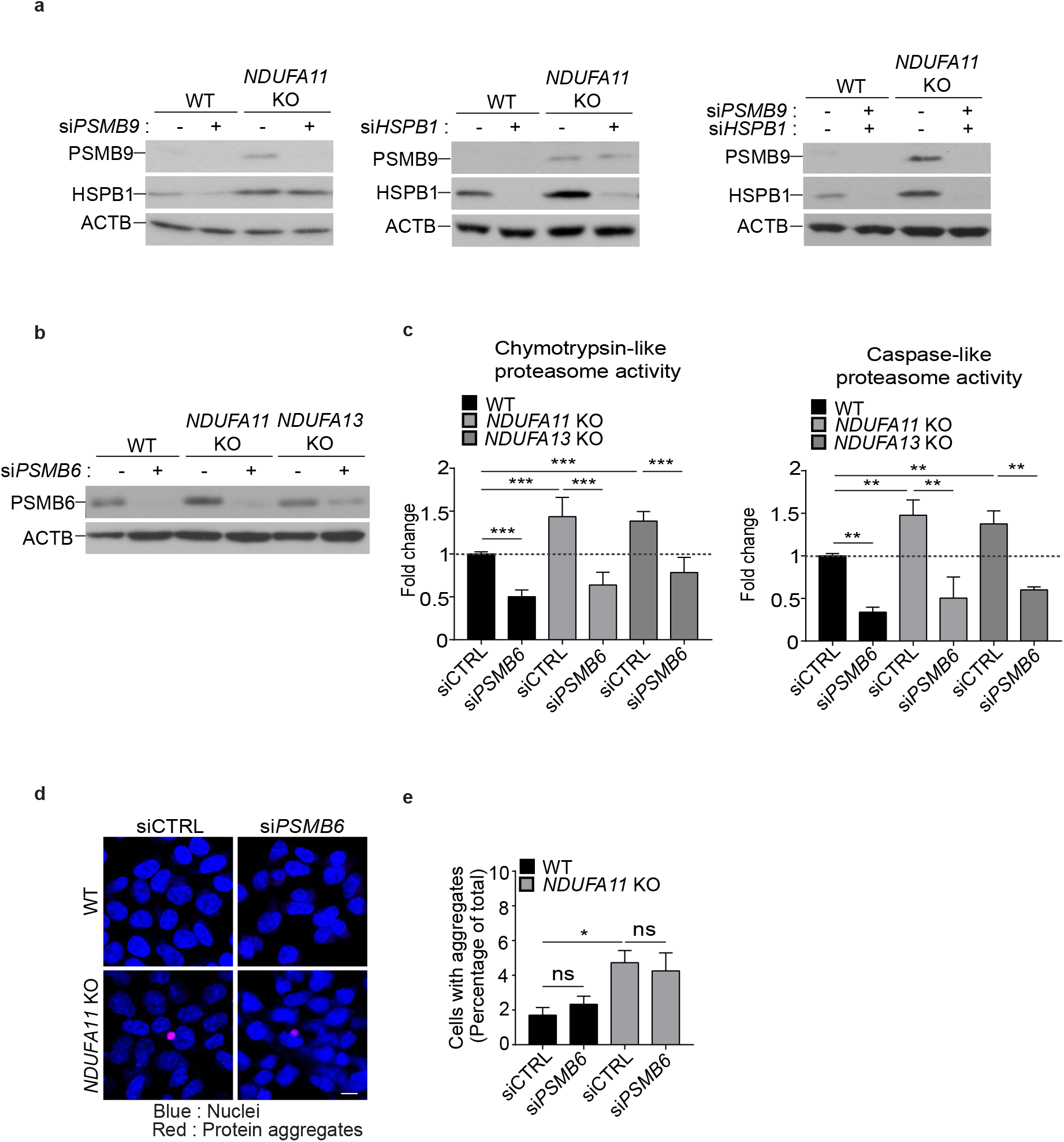
PSMB6 is not responsible to prevent protein aggregates in *NDUFA11* KO HEK293T cells. (**a**) Western blotting validation of knockdown in *NDUFA11* KO and WT HEK293T cells transfected with *PSMB9*, *HSPB1* or control siRNAs for 72 h. (**b**-**e**) Mitochondrial complex I-deficient and WT HEK-293T cells were transfected with *PSMB6* or control siRNAs for 72 h. (**b**) Western blotting validation of *PSMB6* knockdown. (**c**) Chymotrypsin-like and caspase-like proteasome activities in cell lysates presented as fold changes relative to WT. Data shown are mean ± SD. ** p < 0.01, *** p < 0.001; ns, not significant from an ordinary one-way ANOVA with Tukey’s multiple comparisons test or Kruskal-Wallis test with Dunn’s multiple comparisons test using GraphPad Prism. (**d**) Images of protein aggregates stained with ProteoStat. The scale bar represents 10 μm. (**e**) Quantification of the percentage of cells containing aggregates in (**d**). Data shown are representative of 3 independent experiments. Data shown are mean ± SEM. *p < 0.05; ns, not significant from an ordinary one-way ANOVA with Tukey’s multiple comparisons test using GraphPad Prism.

